# Control of cellular cortical tension and shape by RhoGTPase signalling

**DOI:** 10.64898/2025.12.15.694413

**Authors:** Pierre Bohec, Diana Khoromskaia, Manasi Kelkar, Emma Ferber, Guillaume Duprez, Geneviève Lavoie, Léo Valon, Philippe P. Roux, Guillaume Salbreux, Guillaume Charras

## Abstract

Shape changes are ubiquitous in biology, from cytokinesis at the single cell scale to tissue-scale morphogenesis involving coordinated changes in hundreds of cells. In all cases, morphogenesis is powered by gradients in mechanical tension that arise downstream of signalling. Many pathways converge on RhoGTPases that modulate the cytoskeleton and cell contractility to control cell mechanics and, subsequently, shape. Despite their physiological importance, we lack a quantitative understanding of how changes in signalling alter cortical mechanics to drive cell shape change. Here, we use optogenetics to quantitatively characterise the relationship between the amount of RhoGEF localised to the membrane, the downstream myosin recruitment, and the subsequent mechanical changes. We then show that cortical myosin amount and cortical tension increase linearly with the amount of membranous RhoGEF signalling. Based on these data, we develop a predictive model of the temporal evolution of RhoGEF membrane localisation, cortical myosin enrichment, and cortical tension in response to a pulse of light. Using this model together with an active surface model of the cell cortex, we show that the cellular shape changes induced by localised optogenetic recruitment of RhoGEF signalling can be predicted, directly linking gradients in signalling to shape change.

**Significance statement:** Shape changes are ubiquitous in biology, during division in single cells and in tissue during embryogenesis. In all cases, shape change is powered by gradients in mechanical tension that arise downstream of changes in biochemical signals. Despite their importance, we lack a quantitative understanding of how changes in signals alter cell mechanics to drive cell shape change. Here, we control the location and amount of biochemical signal using light to quantitatively characterise the relationship between signals and their resulting biological and mechanical changes. We show that mechanical change scales linearly with the amount of biochemical signal. Based on this, we develop a mathematical model that predicts cell mechanical and shape changes from the location and amount of biochemical signals.

## Introduction

One of the most striking features of living cells is their ability to change shape. As they enter mitosis, cells round up then elongate in anaphase, and generate a cleavage furrow in telophase. These shape changes are controlled by the emergence of tension gradients in the submembranous actin cortex, a thin meshwork of actin filaments, myosin motors and actin-binding proteins ^1,2^. During mitosis, shape change is orchestrated by a complex interplay between the RhoGTPases RhoA and cdc42^3,4^. However, how RhoGTPase activity controls cortical tension to drive shape change remains poorly understood. Yet, precise spatiotemporal control of surface tensions is required for accurate division, while misregulation of cortical contractility can lead to aneuploidy^5^, a factor thought to contribute to cancer progression.

RhoGTPase activity is controlled by RhoGEFs, which activate them by catalysing the exchange of GDP for GTP, and RhoGAPs, which inactivate them by activating hydrolysis of GTP. RhoGEFs typically consist of a regulatory domain, a localisation domain, and a RhoGTPase binding domain, known as the DH-PH domain^6^. Recruitment of a RhoGEF to the cortex leads to changes in RhoGTPase activity that set a complex signalling cascade in motion. This drives changes in cytoskeletal organisation and myosin contractility that ultimately result in mechanical changes and subsequently shape change. Activation of RhoA remodels the F-actin cytoskeleton by stimulating the activity of formins (such as mDia1) and by repressing F-actin severing by cofilin. Simultaneously, RhoA-GTP enhances myosin contraction through the intermediary of Rho-kinase, which phosphorylates myosin light chain to activate contractility and inactivates myosin phosphatase. Together, cytoskeletal remodelling and myosin contractility increase cell surface tension. Previous work has shown that depletion of RhoGTPases changes cell cortical tension^7^ and conversely that increase in RhoA-activity during cell division leads to an increase in cortical tension^8-10^. However, we have little quantitative understanding of how the spatiotemporal recruitment of a RhoGEF to the membrane leads to changes in cortical tension because of a lack of tools to finely tune signalling activity.

Optogenetics has emerged as a key technique to study RhoGTPase signalling, offering tight control on the onset and strength of signalling. Optogenetic actuators can control RhoGTPase activity to modulate cell shape, leading for example to the emergence of lamellipodia in response to Rac1 activation ^11^, cell retraction in response to cdc42 activation^12^, and stress fibre contraction or furrow ingression when RhoA is activated^13,14^. Many optogenetic systems consist of a heterodimeriser pair that associates in response to light. Fine spatial and temporal control is possible by tuning the location, intensity and duration of light exposure^12,13,15,16^. When applied to control of RhoA, these approaches have revealed that cell-generated stresses increase with activation^13,15,17-21^. However, a quantitative understanding of the relationship between the amount of signalling and force generation still remains elusive. This is because theoretical models have so far mostly focused on understanding the spatial and temporal response of the optogenetic heterodimeriser pair to light ^16,22^.

Here, we use optogenetic actuators to relocalise the DH-PH domain of RhoGEFs to the membrane and we monitor the induced mechanical changes using AFM. Relocalisation of RhoGEF to the membrane by a single pulse of blue light leads to recruitment of myosin and an increase in cortical tension with a delay of ∼50s. To separate the dynamics of signalling from those of the optogenetic system, we develop a model of the localisation dynamics of the optogenetic proteins following a single light pulse. This is then combined with a delay model to predict the kinetics and strength of downstream signalling. Using this approach to analyse experiments, we show that the extent of cortical myosin recruitment and the magnitude of increase in cortical tension scale linearly with the amount of RhoGEF relocalised to the membrane. Our framework allows prediction of cellular shape change directly from the spatiotemporal localisation of signalling and paves the way for a quantitative understanding of how signalling controls mechanics and shape.

### Results

### Optogenetic relocalisation of a RhoGEF DH-PH domain to the membrane induces cortical myosin recruitment

To probe the cytoskeletal and mechanical changes induced by RhoGTPase activation, we used a previously established actuator based on light-induced dimerization of the cryptochrome CRY2 with the CIB1 protein^23^. The DH-PH domain of a RhoA specific GEF, p115-RhoGEF/ARHGEF1, is fused to CRY2-mCherry (hereafter CRY2-p115RhoGEF) and CIBN-GFP is targeted to the plasma membrane by fusion to a CAAX domain **(Fig1a)** ^17,24^. In response to blue light, CRY2 changes conformation allowing it to bind CIBN and this leads to relocalisation of the DH-PH domain to the plasma membrane where it can activate RhoA. DH-PH relocalisation thus represents a biological input signal to the RhoA signalling cascade.

For our study, we placed ourselves in a simplified geometry by examining rounded interphase cells to allow robust quantification of relocalisation of signalling to the cell membrane and, simultaneously, precise measurement of the evolution of cortical tension using atomic force microscopy (AFM). As a first step, we examined the biological and mechanical response of cells to a short pulse of light, because in principle this knowledge should allow prediction of the response to an arbitrary light input signal.

In experiments, we illuminate a small region within the cytoplasm of a rounded interphase cell with 488nm light, monitor changes in localisation of CRY2-p115RhoGEF, and quantify the kinetics of relocalisation with an image analysis pipeline (**Supplementary information, Supplementary Fig S1-2**). Prior to activation, CRY2-p115RhoGEF is cytoplasmic but a single pulse of light lasting 250ms leads to relocalisation of ∼10-30% of the total CRY2-p115RhoGEF to the membrane, reaching a peak ∼40s after the pulse (**Fig 1b**). Then, as CRY2 spontaneously reverts to its inactive state and detaches from CIBN, the proportion of membranous CRY2 returns to its original level over a duration of ∼300s, consistent with previous reports^23^. Thus, our control input signal, a single pulse of blue light, gives rise to a biological input signal in the form of relocalisation of a RhoGEF DH-PH domain to the cell membrane.

**Figure 1.**
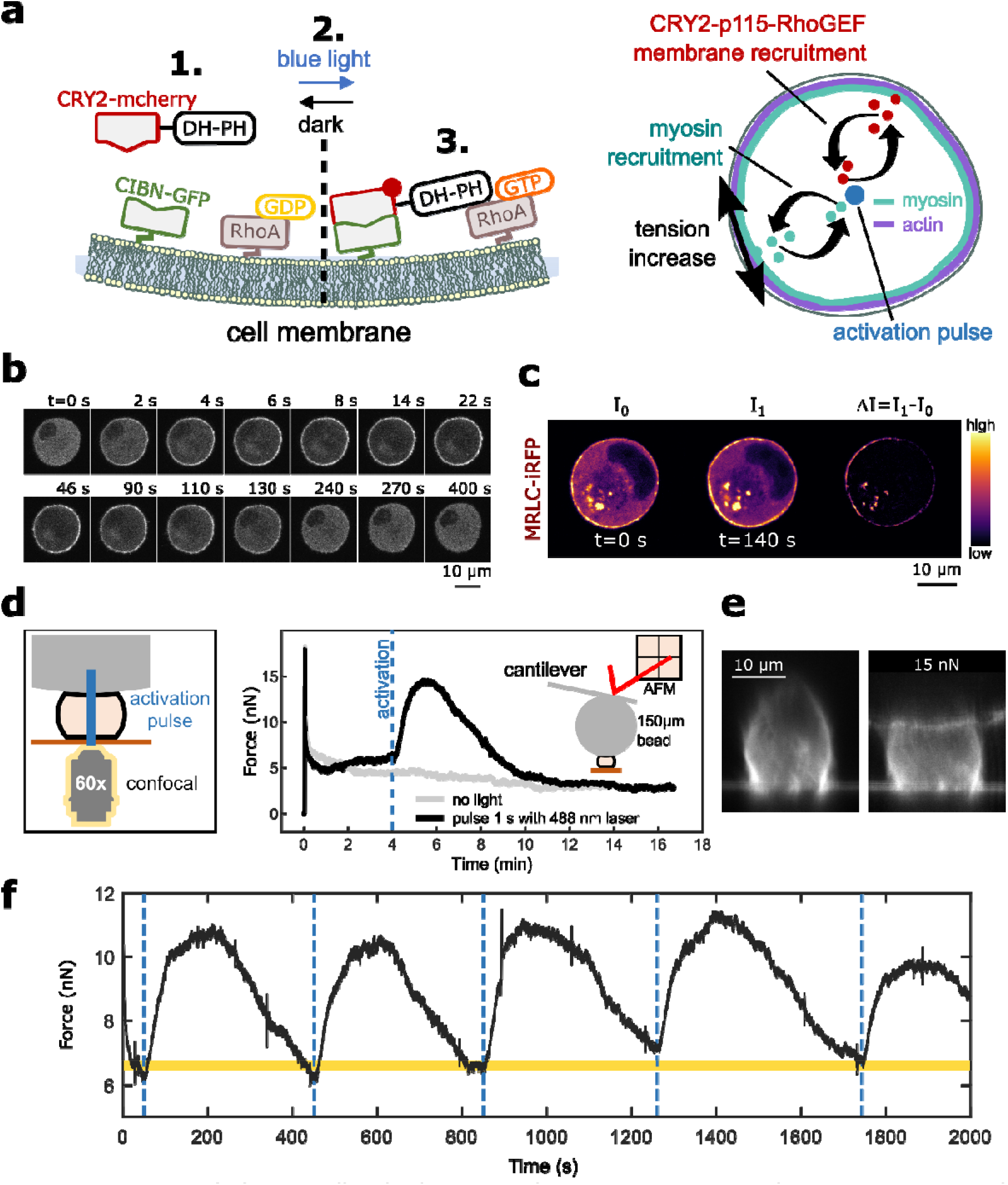
Activation of RhoA signalling leads to cortical myosin recruitment and an increase in cortical tension. **a.** Schematic of the optogenetic actuator. 1. Left panel: In the dark (left), CRY2-p115RhoGEF diffuses freely in the cytoplasm. 2. Exposure to blue light (right), leads to a conformational change in CRY2 enabling it to bind to CIBN at the plasma membrane. This change in localisation brings the DH-PH domain into close proximity with RhoA-GDP and it can catalyse its activation. 3. Right panel: Recruitment of CRY2-p115RhoGEF (red) leads to recruitment of myosin to the cortex (turquoise) and an increase in cortical tension. **b**. Time series of CRY2-p115RhoGEF localisation in a rounded interphase cell. The cell is exposed to a single 250ms pulse of blue light in a small region in its centre at t=0s. **c**. CRY2-p115RhoGEF localisation to the membrane leads to recruitment of myosin to the cortex. The left column shows localisation of MRLC-iRFP before optogenetic activation. The middle column shows localisation after activation. The right column is the difference in fluorescence intensity between the pre and post activation images. Hot colours indicate high intensity and cold colours low intensity. **d**. Cortical tension change in response to optogenetic activation. Left: schematic diagram of the experiment. An AFM cantilever functionalised with a large latex bead (150µm diameter) is brought into contact with a rounded cell. After reaching a threshold force of 15nN, the bead height is kept constant and the force is left to relax for 4 minutes to reach a plateau. Then, a pulse of blue light activates the optogenetic system. Right: Temporal evolution of force measured by AFM in response to compression of a cell exposed to a single pulse of blue light (black curve) and the same cell with no light stimulation (grey curve). Timing of the pulse is indicated by the dashed blue line and only occurs for the black curve. e. Profile view of a cell expressing CAAX-CIBN-GFP before and during compression. **f**. Representative temporal evolution of force in response to multiple pulses of blue light. Actuation pulses are indicated by the dashed blue lines. The initial baseline value for the restoring force is indicated by the yellow shaded region.

As our previous work confirmed that CRY2-p115RhoGEF relocalisation leads to RhoA activation^24^, we examined signalling downstream of RhoA-GTP. As active RhoA modulates the F-actin cytoskeleton and myosin activity, we used cell lines stably expressing our optogenetic actuator alongside reporter constructs for F-actin (LifeAct-iRFP ^24,25^) and myosin (MLC2-iRFP, ^24^). After optogenetic actuation, we could not detect any change in F-actin^24^, perhaps because we use a single pulse of activation lasting 250ms rather than exposure lasting tens of minutes as in previous reports ^13,18^. In contrast, clear myosin recruitment is observed at the cortex (**Fig 1c**). Thus, relocalisation of RhoGEF DH-PH to the membrane provides a well-controlled biological input that can set in motion a signalling cascade that leads to a biological output in the form of myosin recruitment to the cortex.

### Activation of RhoA signalling increases cortical tension

Having established that RhoGEF DH-PH relocalisation induces myosin recruitment to the cortex, we investigated the subsequent changes in mechanics. For this, we compressed rounded interphase cells between a low adhesion coverslip and a large latex bead (∼150μm diameter) glued to an atomic force microscope (AFM) cantilever (**Supplementary Figure S4-5**). In our experiments, we applied a force of 15 nN resulting in a deformation of ∼3.7μm or 25% of cell height and then kept the cell deformation constant (**Fig 1d**,**e**). Within 30s of the application of compression, the restoring force *F*(*t*) relaxes to a plateau, likely due to turn over of actomyosin^26^. In a cortical shell-liquid core model of cell mechanics, the restoring force is proportional to the cell cortical tension *T*(*t*) ^27^ (**Fig 1d**, grey curve; **Supplementary Figure S5**). After the cell reached a mechanical equilibrium, we induced relocalisation of the DH-PH domain to the membrane with a single pulse of blue light and monitored changes in the restoring force for a period of at least 500s. The restoring force *F*(*t*) rose shortly after the light pulse reaching a peak approximately 2-fold larger than the initial force *F*(0) after ∼100s (**Fig 1d, black curve**). *F*(*t*) then decreased over the next 300s, returning approximately to its initial value *F*(0) (**Fig 1d, black curve**). Return to tensional homeostasis followed a time course similar to the loss of membranous localisation of CRY2-p115RhoGEF (**Fig 1b**). Within the same cell, actuation could be repeated multiple times. We stimulated cells at intervals of ∼500s to ensure that CRY2-p115RhoGEF had returned to the cytoplasm after each actuation. In these conditions, each pulse elicited increases in restoring force with similar amplitudes and duration (**Fig 1f**). Remarkably, *F*(*t*) always returned to the same baseline value and the cell showed no sign of plasticity or adaptation, in contrast to experiments in which activation lasts tens of minutes^13,18^.

Overall, our experiments demonstrated that activation of RhoA signalling induced clear changes in cell cortical tension. In the following, we sought to establish a quantitative dynamic relationship between binding of the optogenetic actuator at the membrane, myosin recruitment at the cell surface and cortical tension.

### A temporal delay model can predict recruitment of downstream effectors of RhoGEFs and mechanical changes from the kinetics of the optogenetic actuator

To link RhoGTPase signalling to increase in surface tension, we carried out experiments in which we trigger membrane relocalisation of CRY2-p115RhoGEF with a pulse of blue light and monitor the recruitment of myosin to the cortex using imaging as well as the evolution of cortical tension, which can be calculated from the restoring force measured by AFM^27^.

Using custom written image analysis programmes (**Supplementary information**), we extracted the temporal evolution of membranous CRY2-p115RhoGEF and cortical myosin (**Fig 2a-b**). These measurements showed that, in response to a pulse of light, membranous CRY2-p115RhoGEF, cortical myosin, and cortical tension all displayed very similar temporal evolutions (**Fig 2a-c**). All increased soon after optogenetic activation, reached a peak, and then decayed over a duration of ∼300s. The timing of the peak in myosin and tension appeared delayed compared to the peak in CRY2-p115RhoGEF, as expected given they are downstream of the activation of RhoA by CRY2-p115RhoGEF. We then verified the role of myosin in the rise in cortical tension. For this, we first stimulated cells in control conditions and measured their increase in cortical tension before repeating actuation in the presence of a photostable derivative of the myosin inhibitor blebbistatin (s-nitro-blebbistatin) (**Fig 2d**). Myosin inhibition abrogated the rise in cortical tension, confirming a causal link between myosin recruitment and the rise in tension (**Fig 2e-f**).

**Figure 2.**
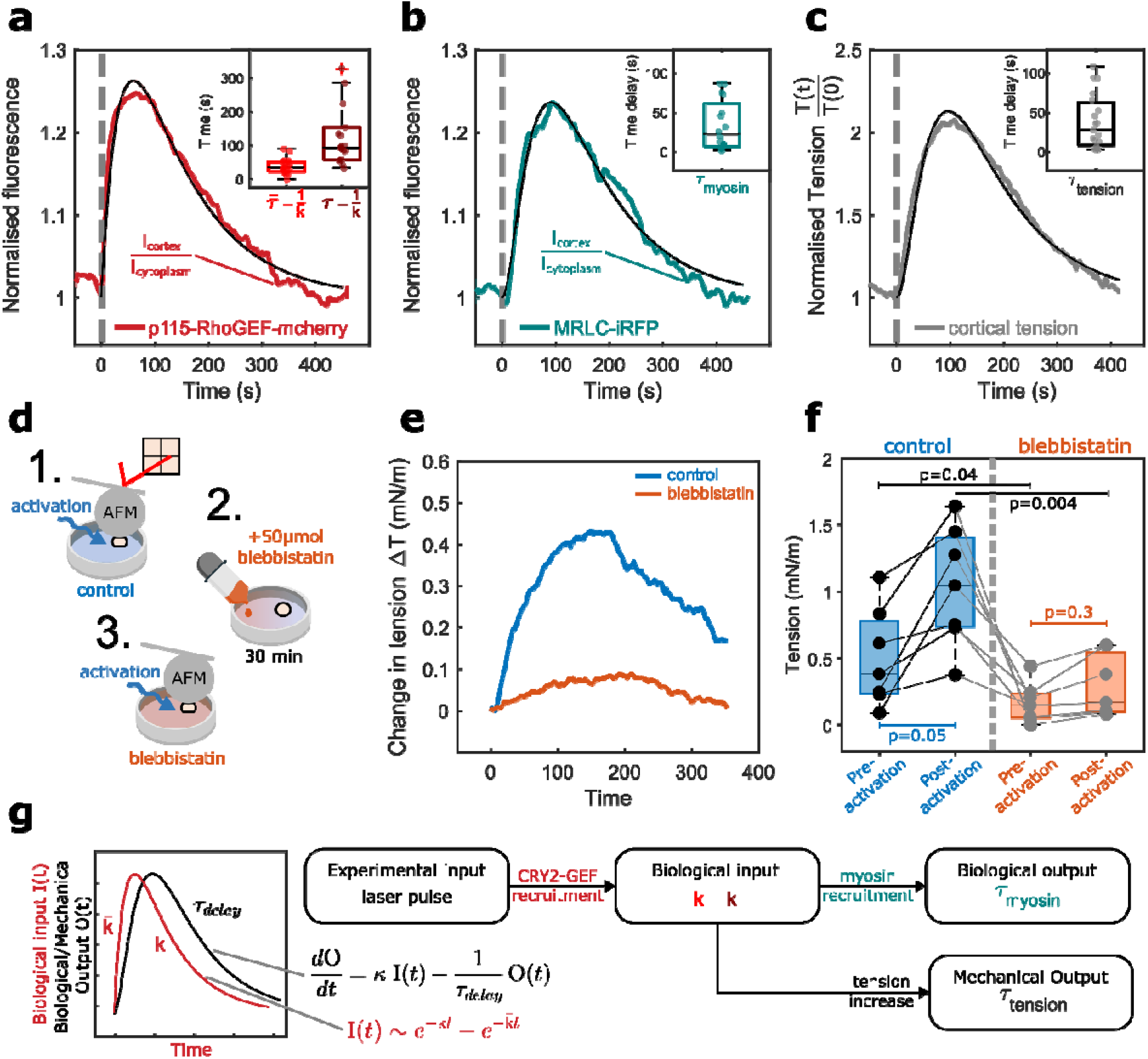
The kinetics of myosin recruitment and tension increase can be predicted with a reaction and delay model. **(a-c, f)** Boxplots indicate the 25th and 75th percentiles, the black line within the box indicates the median, and the whiskers extend to the most extreme data points that are not outliers. Outliers are indicated by red crosses. Individual data points are indicated by solid dots. Wilcoxon rank-sum test p-values are displayed on the graph. **a.** Temporal evolution of normalised fluorescence of CRY2-p115RhoGEF in response to a single pulse of blue light (n=22 cells). Actuation is indicated by the dashed grey line. Membrane fluorescence intensity is normalised to the fluorescence intensity in the cytoplasm. Solid red line indicates the mean and the shaded area the standard error. Solid black line shows the fit to a function of the form . Inset shows the characteristic times and . b. Temporal evolution of normalised MRLC-iRFP fluorescence intensity in response to a single pulse of blue light (n=16 cells). Actuation is indicated by the dashed grey line. Solid blue line indicates the mean and the shaded area the standard error. Solid black line shows the fit to equation 1. Inset shows the characteristic myosin delay . **(a-b)** Curves display normalised to the value at time 0: . **c**. Temporal evolution of normalised cortical tension in response to a single pulse of blue light (n=19 cells). Actuation is indicated by the dashed grey line. Solid black line shows the fit to equation 1. Inset shows the characteristic tension delay . Tension is normalised to its value at time 0. d. Schematic of the experiment shown in **(e-f)**. First, the response of a cell to optogenetic stimulation is characterised as in Fig 1d. Then S-nitro-blebbistatin is added to the medium at a concentration of 50μM. Optogenetic actuation is then repeated on the same cell in the presence of blebbistatin. **e**. Temporal evolution of the change in cortical tension Δ *T*(*t*)= *T*(*t*) − *T*(0) following exposure to a pulse of blue light in control conditions (blue line) and in the presence of blebbistatin (orange line). The solid line indicates the mean and the shaded area denotes the standard error. Average of 7 experiments. f. Cortical tension before and after activation in control conditions and in the presence of blebbistatin. Solid lines link data points from the same cell. **g**. Principle of the delay model. Left: Temporal evolution of the biological input I(t) in red and an output signal O(t) in black. Right: experimental input is a pulse of blue light. This gives rise to recruitment of CRY2-p115RhoGEF to the cell membrane (characterised by the fluorescence intensity I(t)) with kinetics that can be fitted with a sum of two exponentials with parameters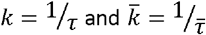 . Membranous CRY2-p115RhoGEF represents a biological input signal that gives rise to a biological output (e.g. recruitment of myosin) with a delay *τ*_*myosin*_ and a mechanical output, the change in tension, with a delay *τ*_*tension*_.

When we compared the temporal evolution of cortical myosin and cortical tension, we noticed they had similar kinetics to the temporal evolution of membranous CRY2-p115RhoGEF except for a delay in reaching their peak. As prediction of the response directly from biochemical reactions is not possible due to large uncertainty on many molecular variables, we adopted a coarse-grained approach that predicts the evolution of cortical myosin and tension by assuming that any given step of the signalling cascade can be predicted by rescaling of the input signal and adding a temporal delay that becomes progressively longer with the number of steps in the signalling cascade^28^. In our experiments, the relevant biological input is the surface concentration of membranous CRY2-p115RhoGEF, [CRY2^*^ - CIBN]^2D^ (*t*) (with CRY2* denoting activated CRY2, **Fig 2a**). Based on empirical evidence from previous studies^12^ and the shape of the temporal evolution of membrane localised CRY2-p115RhoGEF, we described its temporal evolution with a function of the form 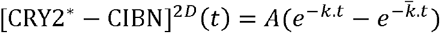 and verified that this fitted experimental data of cortical fluorescence intensity well (black line, **Fig 2a**). Knowing this input, a downstream output *O*(*t*) that responds with a simple temporal delay *τo* will obey the following equation (**Fig. 2g, Supplementary information**):

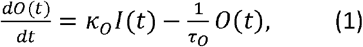

where *κ*_*o*_ is a rescaling factor whose units depend on the nature of the output *O*(*t*), *τ*_*o*_ is a delay that is specific to *O*(*t*), and I(t).

In practice, we first find *k* and 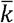 by fitting the dynamics of normalized surface concentration of CRY2 estimated by Δ*I*_*cortex*_ (*t*)/Δ*I*_*cyto*_ (*t*), the average ratio of intensities above background of CRY2-p115RhoGEF in the membrane relative to cytoplasm (see **Supplementary information**). In our experiments, 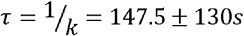 was several fold larger than 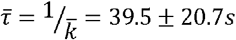, signifying that the long time deactivation occurs on the characteristic time τ (**Fig 2a**, inset). Then, we fitted the temporal evolution of myosin recruitment and cortical tension with equation (1) with k and 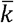 as fixed parameters to obtain τ_o_. After fitting, the predicted curves agreed remarkably well with the experimental data (black lines, **Fig 2b-c**). Myosin recruitment occurred with a delay *τ*_*myosin*_∼33.8 ± 31.6s compared to the timing of CRY2-p115RhoGEF relocalisation to the membrane and tension increase occurred with a delay of *τ*_*ension*_ =38.4 ± 34.4S. These delays reveal the duration necessary to set in motion a complex signalling cascade downstream of RhoGEF recruitment, indicating a lower bound for the rate at which cellular morphogenetic changes can take place in response to a change in signalling. Interestingly, the delay between CRY2 relocalisation to the membrane and myosin recruitment is of a similar magnitude to the delay measured between RhoA activation and myosin recruitment in contractile actomyosin foci in the *C. elegans zygote* (∼20s, ^29^).

In summary, this coarse-grained approach uses the temporal evolution of signalling controlled by optogenetics to predict the temporal evolution of its downstream outputs in the form of myosin recruitment to the cortex and cortical tension.

### DH-PH domains from different RhoGEFs lead to distinct mechanical outputs

The RhoGEF family comprises many proteins^30^ and previous work has shown that a same RhoGEF can induce distinct responses dependent on context because some RhoGEFs can interact with different RhoGTPases^31^. However, we do not know if different RhoA-specific RhoGEFs induce distinct mechanical responses. We investigated whether different RhoGEF GTPase binding domains can induce distinct mechanical responses. For this, we examined the mechanical response elicited by two closely related RhoGEFs (p115RhoGEF and PDZRhoGEF) and one more distant one (Ect2)^6^. We generated actuators based on their DH-PH domains (CRY2-mCherry-PDZRhoGEF-DH-PH and CRY2-mCherry-Ect2-DH-PH termed CRY2-PDZRhoGEF and CRY2-Ect2 hereafter, **Fig 3b-c**) and characterised the cortical tension response to a single pulse of blue light using the same protocol as previously (**Fig 3a**).

**Figure 3.**
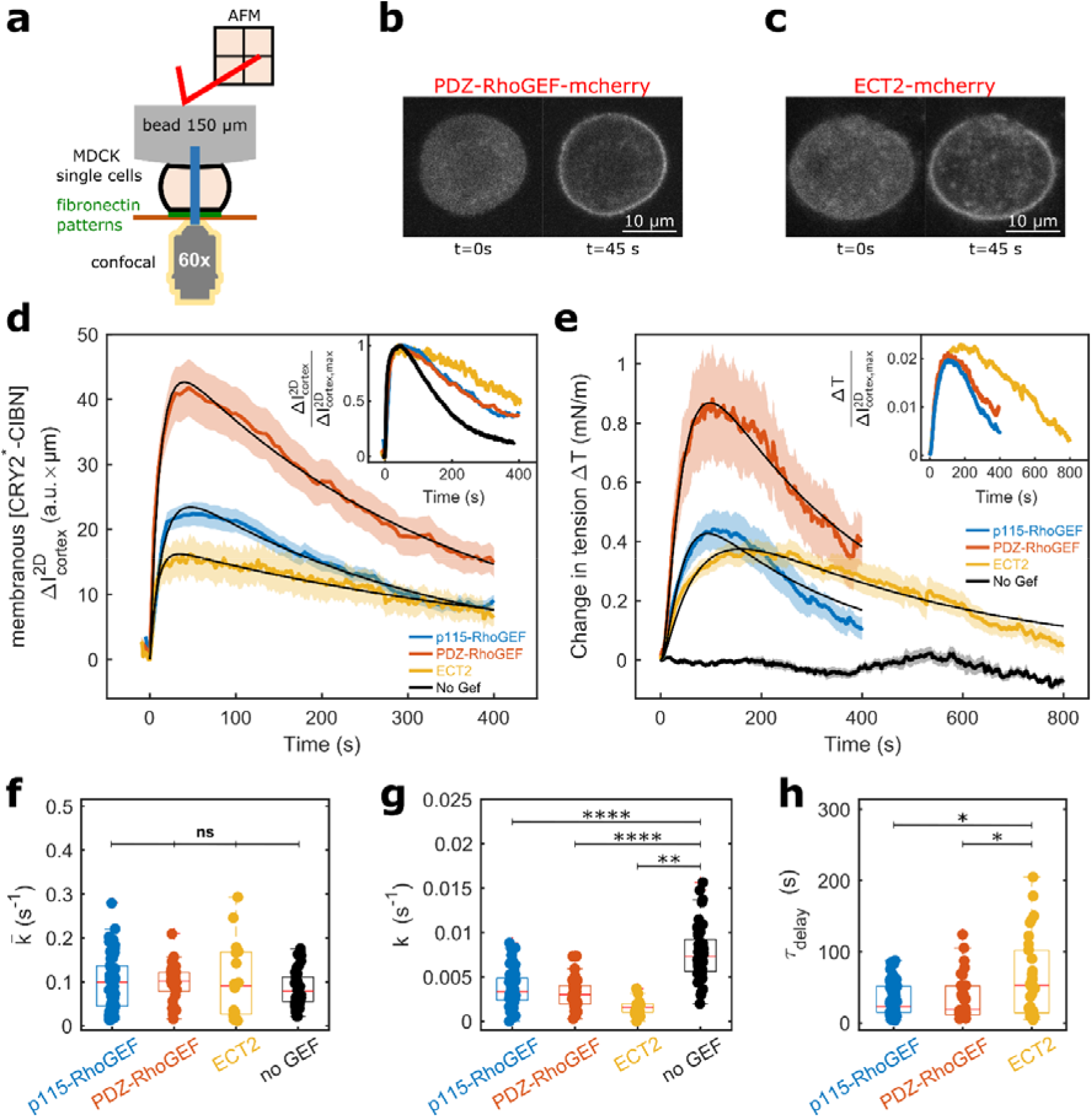
DH-PH domains from different RhoGEFs lead to distinct mechanical outputs. **a.** Schematic diagram of the experiment. Rounded MDCK cells seeded on fibronectin micropatterns are squeezed under a 150µm latex bead attached to an AFM cantilever and are stimulated by a 250ms pulse of blue light. **(b-c)**. Representative localisation of CRY2-PDZRhoGEF (b) and CRY2-Ect2 (c) before and after a blue light pulse. **d**. Temporal evolution of membranous [CRY2*-CIBN], as measured with the cortical fluorescence intensity (Supplementary Information), following illumination. Cells were exposed to a pulse of blue light lasting 250ms at t=0s. Inset shows the temporal evolution normalised to its maximum. The CRY2-mCherry with no GEF domain is not shown on the main graph because its expression is many fold higher than the CRY2-mCherry fused to RhoGEF domains (Fig S6). The number of cells examined is n=64 for p115RhoGEF, n=33 for PDZ-RhoGEF, n=17 for Ect2 and n=45 for CRY2-mCherry with no GEF domain. Solid lines indicate the mean and the shaded area shows the standard error. Black lines show a fit to a function of the form . **e**. Temporal evolution of the change in cortical tension in response to a pulse of blue light. Light stimulation occurred at t=0s. Inset shows the change in tension normalised to the maximum cortical fluorescence intensity of CRY2*-CIBN . The number of cells examined is n=61 for p115RhoGEF, n=35 for PDZ-RhoGEF, n=33 for Ect2 and n=15 for CRY2-mCherry with no GEF domain. **(f-h)** Distribution of,, and for different RhoGEF domains. The number of cells examined is n=61 for CRY2-p115RhoGEF, n=35 for CRY2-PDZ-RhoGEF, n=33 for CRY2-Ect2 and n=15 for CRY2-mCherry with no GEF domain. Boxplots indicate the 25th and 75^th^ percentiles, the red line within the box indicates the median, and the whiskers extend to the most extreme data points that are not outliers. Outliers are indicated by red crosses. Individual data points are indicated by solid dots. Wilcoxon-rank sum test, n.s.: p>0.05, *: p<0.05, **: p<0.01, ***: p<0.001, ****: p<0.0001. (a-e) Individual curves and their fits are shown in Supplementary Fig S5.

We found that all CRY2-RhoGEFs relocalised to the membrane with similar kinetics (**Fig 3d**), likely because this is dominated by binding of activated CRY2 (CRY2*) to CIBN. The signal then returned towards its baseline as CRY2* inactivated. In all cases, membrane accumulation of CRY2* was followed by an increase in cortical tension with kinetics that appeared to follow those of the CRY2 signal (**Fig 3e, Supplementary Fig S6**). Intriguingly, the maximum average change in tension was different between GEF domains, with CRY2-PDZRhoGEF giving the largest increase followed by CRY2-p115RhoGEF and CRY2-Ect2 (**Fig 3e**). We reasoned that this could be due to differences in the expression of signalling proteins (CRY2-RhoGEF) or downstream effectors (RhoA or myosin MYH9 and MYH10). When we compared the abundance of mRNA transcripts for these proteins across cell lines, we found no difference in RhoA and MYH10, and only a small difference in MYH9 (**Supplementary Fig S7b**). In contrast, there was a large difference in mRNA transcripts levels for the different CRY2-RhoGEFs with CRY2-PDZRhoGEF being the most abundant followed by CRY2-p115RhoGEF (**Supplementary Fig S7b**). Therefore, we hypothesised that differences in the maximum amplitude of tension change were likely due to differences in the abundance of actuators across cell lines. To test this hypothesis, we normalised the tension change to the maximum concentration of CRY2 at the membrane determined from the fluorescence signal. This revealed that the amplitude of the change in cortical tension per unit CRY2-RhoGEF fluorescence was similar for all RhoGEFs (inset, **Fig 3e**).

To quantify these data, we fitted the experimental curves as previously. 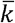 was similar for all conditions (**Fig 3f**), whereas *k* was an order of magnitude smaller. Interestingly, *k* was significantly smaller for all CRY2-RhoGEFs than for CRY2 alone (No GEF, **Fig 3g**). As *k* dominates deactivation kinetics, this indicated that detachment from the membrane was slower when a RhoGEF domain was present. The significant difference in unbinding from the membrane compared to CRY2 alone likely arises because the RhoGEF domains bind to partners that stabilise its membrane localisation. Finally, *τ*_*delay*_, which quantifies the kinetics of the tension change induced by each actuator was significantly longer for Ect2 than for PDZRhoGEF and p115RhoGEF (**Fig 3e, 3h**), maybe indicating differences in binding partners or downstream signalling cascade.

Thus, the RhoGEFs that we examined all induced a similar change in cortical tension per actuator protein relocalised to the membrane but appeared to induce tension responses with different kinetics, as evidenced by differences in their delays.

### A kinetic model of optogenetic relocalisation to the membrane

Although the temporal evolution of membrane-bound CRY2 was well-fit by a double exponential, previous work did not reveal the biochemical basis for this functional form^12^. Such knowledge is essential to understand how an arbitrary light input signal translates into a change in signalling, recruits downstream effectors, and changes mechanics.

Therefore, we sought to predict the kinetics of CRY2-mCherry localisation to the membrane based on the biochemical reactions that underlie the response of the optogenetic system to a pulse of blue light (**Fig 1b, 2a**). After exposure to light, a portion of the total CRY2 protein becomes activated to CRY2*, which can bind CIBN at the membrane to form a complex CRY2*-CIBN with a rate constant *k*_+_ (**Fig 4a**). We assume that inactivation of CRY2* follows a first order process with rate constant *k*, that free and bound CRY2* inactivate with the same rate, and that inactivated CRY2 immediately dissociates from CIBN (**Fig 4a**). These assumptions lead to the following simplified reaction scheme for CRY2*:

**Figure 4.**
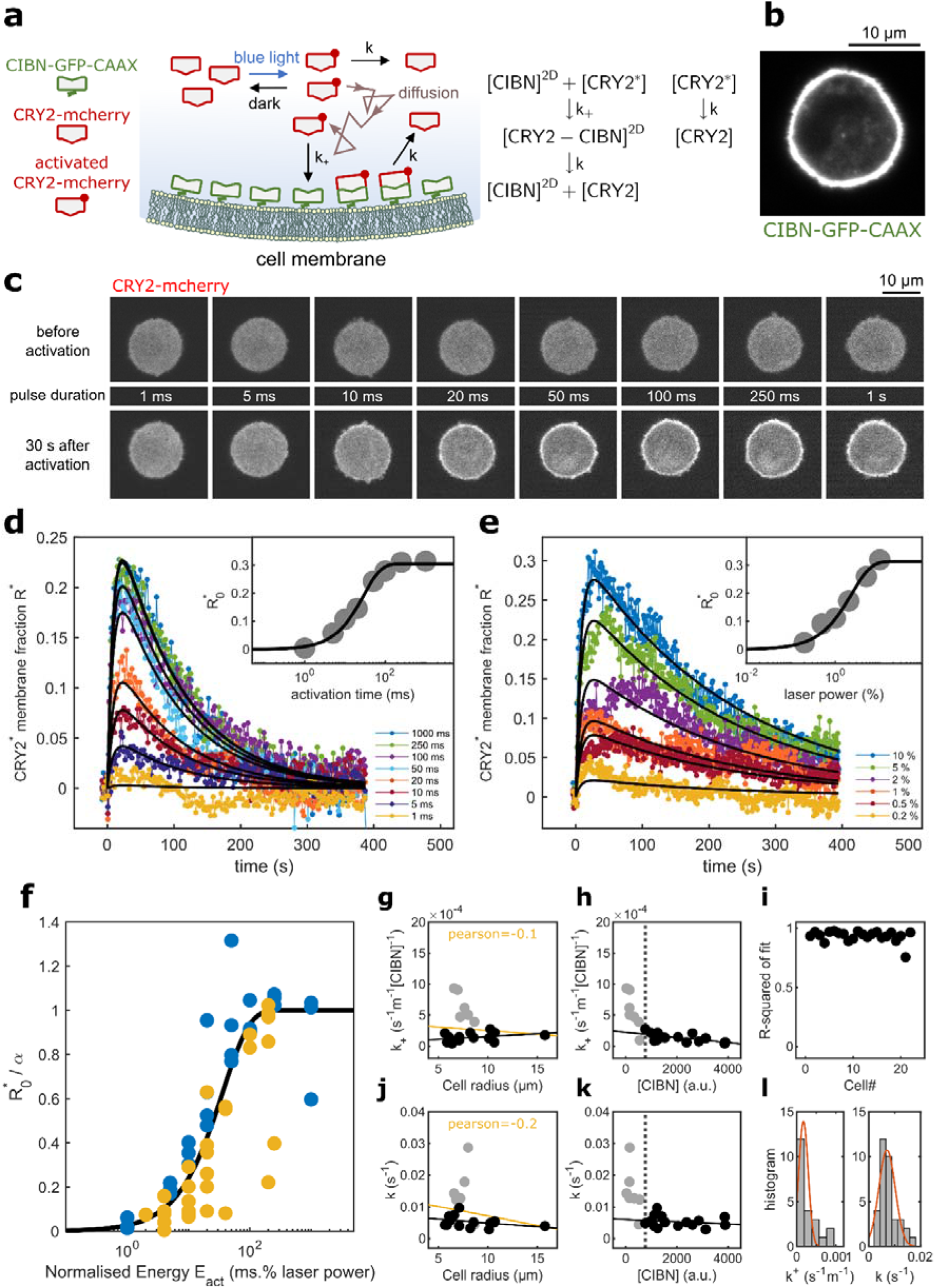
Kinetic model of the optogenetic actuation system. **a.** Left: Schematic diagram of the dynamics of CRY2-mCherry before and after activation. Right: Reaction scheme for CRY2 and CIBN. **b**. Representative localisation of CIBN-GFP-CAAX. **c**. Relocalisation of CRY2-mCherry as a function of the duration of the activation pulse. Top row: CRY2-mCherry localisation prior to exposure. Middle row: pulse duration. Bottom row: CRY2-mCherry localisation 30s after exposure to blue light. All panels show the same cell. Following exposure to light, the cell recovered for 400s to allow full inactivation of CRY2. Scale bar=10µm. **d**. Temporal evolution of the membrane fraction of CRY2 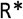 for different durations of light pulses and fixed laser intensity (1%). Solid lines indicate fits with the analytical model. Inset: 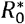 as a function of light pulse duration. **e**. Same plot as (d) for light pulses with varying laser intensity but a fixed duration (20 ms). Solid lines indicate fits with the analytical model. Inset: 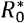 as a function of the laser intensity of the light pulse. **f**. 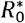 as a function of laser energy divided by the maximum photoconversion efficiency . Experiments in which laser power was varied are plotted in yellow. Experiments in which duration of activation was varied are plotted in blue. The solid black line shows a fit of the form 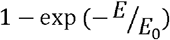 to the data. **g, h, j, k**. Rate constants of binding (*k* +) and unbinding (*k*) between CRY2* and CIBN as a function of cell radius **(g, j)** and as a function of CIBN concentration (h,k). Data points for cells with [CIBN] < 760 a.u. are shown in gray. **i**. Goodness of fit represented by the R-squared values, plotted for each of the 22 cells. l. Distribution of *k*+and . The orange line represents a fit to a normal distribution.

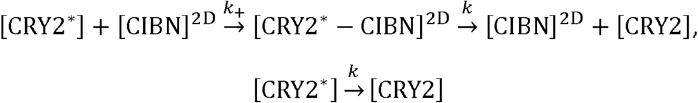

where [CRY2^*^],[CRY2] are the three-dimensional concentrations of active and inactive CRY2 in the cytoplasm, and [CIBN]^2D^,[CRY2^*^ - CIBN]^2D^ the two-dimensional concentrations of CIBN and CRY2*-CIBN complexes at the cell membrane. Using qPCR, we verified that, in all of our optogenetic cell lines, CIBN transcripts are at least 100-fold more abundant than CRY2 transcripts (**Supplementary Fig S7a**), indicating that binding sites at the membrane for CRY2* are far from saturation, such that we take [CIBN] ≃ [CIBN]_tot_, the total CIBN concentration at the cell surface. Solving the chemical kinetic equations, we find that, following a transient pulse of activation, the temporal evolution of CRY2* bound to the membrane [CRY2*-CIBN](t), and CRY2* in the cytoplasm [CRY2*](t), with the initial condition [CRY2^*^ - CIBN]^2D^(*t*= 0) = 0 are given by (**Supplementary theory**):

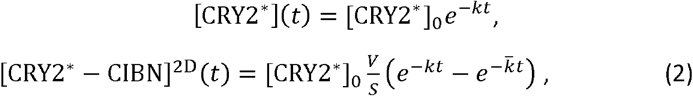

where *V* is the cell volume, *S* the membrane surface area, [CRY2^*^]0 the initial concentration of activated CRY2*, and we have introduced the rates *k* and 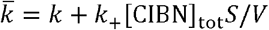. We note that the concentration [CRY2^*^ — CIBN]^2D^ in Eq. (2) has the same functional form as the function chosen to fit *I*(*t*) in **Figs 2-3**: *k* is now reinterpreted as the inactivation rate of CRY2* and 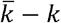 is proportional to the binding rate of CRY2* and CIBN. In our data analysis, we estimated the proportion of CRY2 at the membrane relative to the total CRY2 in the cell, R* (see **Supplementary Information**):

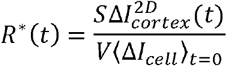

With S the cell surface, V the cell volume, and ⟨ Δ*I*_*cell*_ ⟩ _t=0_ the average intensity of CRY2 above background in the cell at time 0^-^. The temporal evolution of R* follows the equation (see **Supplementary Information**):

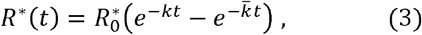

With 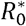 a parameter proportional to [CRY2^*^]0.

### CRY2 relocalisation is proportional to illumination

We next sought to understand the dynamic range of our optogenetic system by quantifying the proportion of activated CRY2 as a function of our experimental control variables: the intensity of the light pulse and its duration.

First, we performed a sweep in pulse duration keeping laser power constant (**Fig 4c, d**). In each cell examined, we exposed a region of 4×4 (0.4×0.4µm) pixels located in the cytoplasm to sequential pulses of 488nm light with durations ranging between 1 and 1000ms (with 1% of the maximal laser power), imaged the cell response at 1s intervals, and computed the membrane fraction of CRY2 R* for each time point (**Methods**). We followed the cells for 400s because CRY2 returns to the cytoplasm within this time frame (**Fig 1b**). The amplitude of R* increased as a function of pulse duration up to ∼250ms, after which it saturated (**Fig 4c, d**). Next, we kept the pulse duration constant (20 ms) but varied laser power from 0.2-10% of the maximal laser power. Again, the amplitude of R* rose with laser power (**Fig 4e**).

Interestingly, the proportion of total CRY2 activated R*_0_ increased both with pulse duration and laser power. This indicates that R*_0_ is controlled by the energy of the light stimulus. When we plotted R*_0_ as a function of the laser energy, we found that the data from both experiments collapse onto the same curve (**Fig 4f**). This suggested that the CRY2 conformational change in response to illumination followed a function of the form:

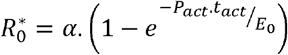

With *α* the maximum photoconversion efficiency, *E*_0_ a characteristic energy of photoconversion, *P*_*act*_ the laser power, and *t*_*act*_ the exposure time. We found *α* = 0.48 ± 0.12, indicating that only ∼48% of total CRY2 is photosensitive. This dependency is consistent with a probability of activation proportional to laser power during the time of activation (see **Supplementary Information**).

We next examined the dependency of *k* and *k*_+_ on the cell size and the level of expression of membranous CIBN. While our cell lines generally have high levels of CIBN transcripts compared to CRY2 (**Supplementary Fig S6a**), they nevertheless display cell to cell variation in CIBN expression. We took advantage of this and plotted *k* and *k*_+_ as a function of CIBN fluorescence intensity. When CIBN fluorescence was sufficiently high (i.e. >760 a.u.), neither rate constant depended on CIBN fluorescence (black dots, **Fig 4h, k**). This confirmed that, in cells with a sufficiently high CIBN expression, binding sites for CRY2 are far from saturated and loss of affinity due to occupation of CIBN molecules can be neglected. Then, we examined dependency of *k* and *k*_+_ on the cell radius. In the cells with high CIBN fluorescence, we found that neither *k* nor *k*_+_ depended on cell radius (black dots, **Fig 4g, j**), indicating that the binding of CRY2* to CIBN is likely not diffusion-limited. Cells with high CIBN expression therefore formed a distinct peak in the distribution of *k* and *k*_+_ (**Fig 4l**).

In summary, our kinetic model allows prediction of the portion of CRY2 localised to the membrane following activation by a pulse of blue light. When CRY2 is fused to a RhoGEF DH-PH domain, equation (2) gives the temporal evolution of the amount of signalling as function of time. This serves as the biological input for equation (1) that predicts the downstream effects of the signalling cascade, such as myosin recruitment to the cortex and cortical tension increase. Thus, together, equations (2) and (1) allow to link a light signal to a change in mechanics.

### Myosin recruitment and cortical tension are proportional to the amount of RhoGEF signalling

Next, we sought to experimentally validate the linearity of the response captured by equation (1), which implies that myosin recruitment and tension change should vary linearly with the amount of membranous RhoGEF. For this, we examined the change in cortical myosin and cortical tension in response to blue light pulses of varying duration.

In our experiments, we probed naturally rounded mitotic cells because changes in RhoA activity and cortical tension are key to driving shape changes leading to cytokinesis. We blocked cells in metaphase with MG-132 because this ensures a stable cortical tension^8^, unlike block with nocodazole which frees GEF-H1 and leads to localised pulses of RhoA activity ^21^.

We subjected each cell to sequential pulses of blue light ranging from 20 to 1000ms and acquired images of CRY2-p115RhoGEF and myosin-iRFP using confocal microscopy while simultaneously measuring the evolution of cortical tension with AFM for 400s (**Fig 5a-c**). We determined the membrane fraction of CRY2 R* and the cortical fraction of total myosin R*_myosin_ at each time point (See eqs (4-5), **Supplementary information**).

**Figure 5.**
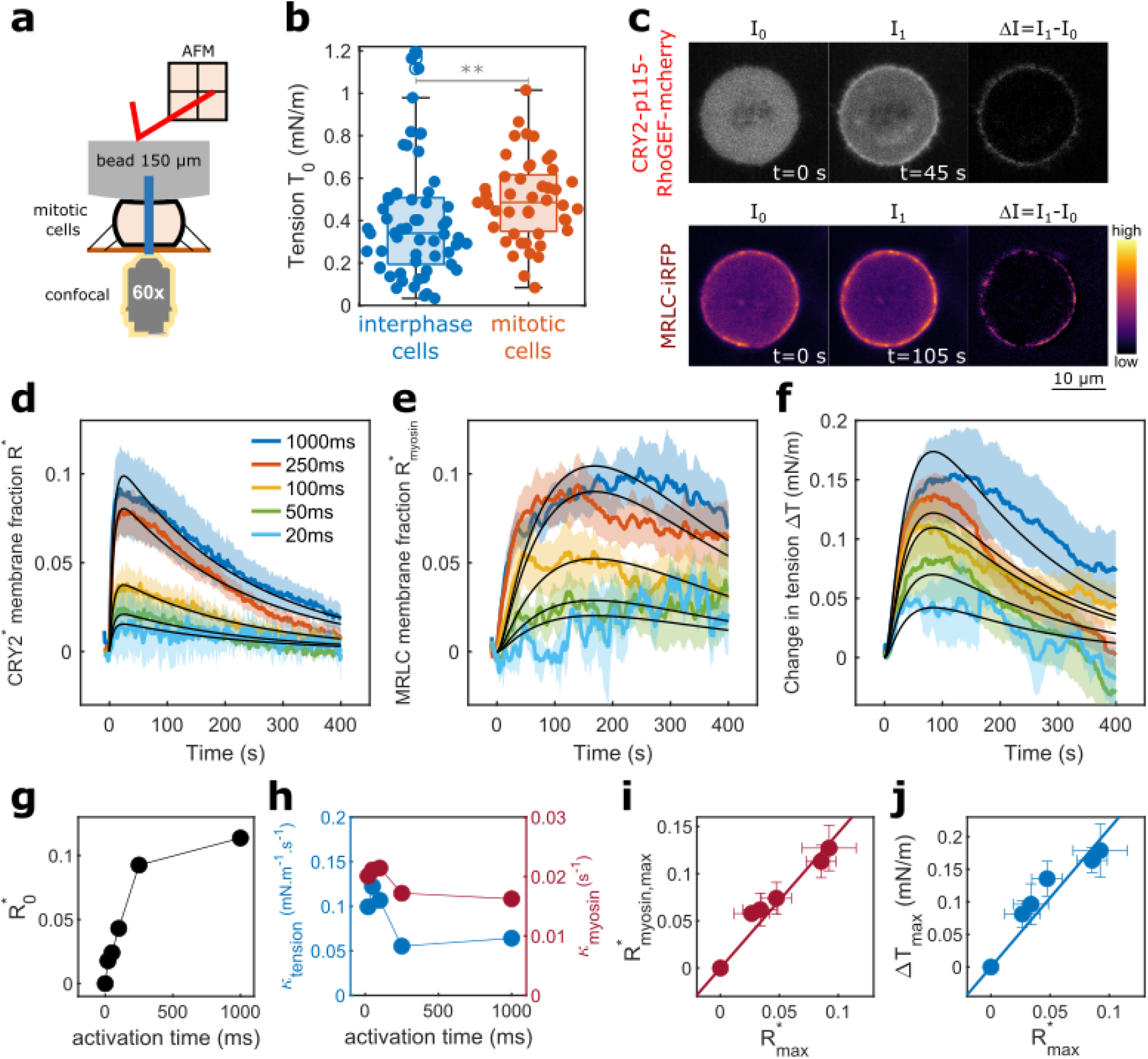
Myosin recruitment and tension increase are proportional to the amount of membranous CRY2-p115RhoGEF. a. Schematic diagram of the experiment. **A** mitotic cell is squeezed between a coverslip and a large latex bead. At t=0s, a pulse of blue light is delivered to a small region in the centre of the cell. Recruitment of CRY2-p115RhoGEF and MRLC-iRFP are monitored by imaging and the change in tension is monitored by AFM. **b. Cortical** tension distribution in interphase cells (n=58) and mitotic cells (n=45) prior to activation. Boxplots indicate the 25th and 75th percentiles, the black line within the box indicates the median, and the whiskers extend to the most extreme data points that are not outliers. Outliers are indicated by red crosses. Individual data points are indicated by solid dots. Wilcoxon rank-sum test, p=0.007. **c.** Change in membranous CRY2-p115RhoGEF (top row) and cortical MRLC-iRFP (bottom row) induced by light exposure at t=0^+^s. Columns show the fluorescence intensity prior to exposure (I_0_), the fluorescence intensity at the peak of recruitment (I_1_), and the change in intensity . Scale bar 10μm. **d**. Temporal evolution of the membrane fraction of CRY2 R* in response to different durations of light pulses. **e**. Temporal evolution of the cortical fraction of MRLC-iRFP R*_myosin_ in response to different durations of light pulses. **f**. Temporal evolution of the change in cortical tension in response to different durations of light pulses. (d-f) For each curve, exposure occurred at t=0s and the signal was followed for 400s. Different durations of activation are indicated by different colours. The solid line indicates the mean and the shaded area shows the standard error. The solid black lines show a fit to equation (2) for d and to equation (1) for e-f. **g**. Proportion of total CRY2 activated, R*_0_, as a function of exposure time obtained from fitting to equation (2). Individual curves are presented in Supplementary Fig S7. **h**. Rescaling factors *κ*_myosin_ (right y-axis, red) and *κ*_tension_ (left y-axis, blue) obtained by fitting Eq. S24 to the averaged R*_myosin_ and tension responses (panels e-f). For these fits, the temporal delay *τ* was taken to be the same for all activation times. The parameters *k* and 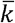 were also shared across all activation times and were obtained independently from fitting the data in d using Eq. 3. **i**. Maximum change in cortical fraction of myosin R*_max,myosin_ as a function of the maximum of the membranous fraction of CRY2 R*_max_. Each data point represents the average for all of the experiments and the whiskers denote the standard error. j. Maximum change in tension Δ _max_ as a function of function of the maximum of the membranous fraction of CRY2 R*_max_. **(i-j)** For the 0 ms activation time, values were taken as the average of the data points during the 31s preceding activation; error bars are not visible because the standard deviations are extremely small. Each data point represents the average for all of the experiments and the whiskers denote the standard error. **(d-i)** The number of cells examined for each duration is n=12 for t_act_=1000ms, n=24 for t_act_=250ms, n=8 for t_act_=100ms, n=8 for t_act_=50ms, n=4 for t_act_=20ms.

As expected, the amplitude of the evolution of R* increased with increasing pulse duration (**Fig 5d**). The change in the proportion of cortical Myosin-iRFP, R*_myosin_, and the increase in cortical tension *ΔT* also increased with increasing pulse duration (**Fig 5e-f**).

We next investigated the relationship between the proportion of membranous RhoGEF, cortical myosin, and the change in cortical tension. The proportion of activated CRY2-p115RhoGEF, quantified with R*_0_, increased as a function of activation time, first increasing before saturating beyond 250ms exposure (**Fig 5g**). For each output from signalling (myosin recruitment and tension), rescaling factors to change (black curves, **Fig 5e-f**). The kinetic parameters *k* and *k*- were kept we fitted the response with Eq S24 using a common delay for all pulse durations but allowing the identical for all activation curves and taken from fitting the data taken on the same cells using equation (3) (**Fig 5d**, black lines). This approach gave delays for the rise in myosin and tension, *τ*_*myosin*_ = 120 and *τ*_*tension*_ = 35*s* and rescaling factors *κ*_*tension*_ = 0.1 ± 0.03 *mN. m*^−1^.*s*^−1^ (**Fig 5h**, blue) and *κ*_*myosin*_ = 0.02 ± 0.002 *s*^-1^ (**Fig 5h**, red). Although the values *κ*_*tension*_ and *κ*_*myosin*_ were fitted separately for each pulse duration, their variation was small or did not show a clear trend as a function of activation time (**Fig 5h**), consistent with an approximately linear relationship between R* and the responses of myosin and tension. To further verify the postulated linear relationship between RhoGEF relocalisation and myosin recruitment or tension, we plotted the maximum proportion of cortical myosin R*_max, myosin_ and the maximum change in tension Δ*T* as a function of the maximum proportion of membranous CRY2-p115RhoGEF, R*_max_. This confirmed a linear relationship in both cases (**Fig 5h-i**). Together these data revealed that, in mitotic cells, myosin recruitment and tension increase are linearly proportional to the amount of membranous CRY2-p115RhoGEF relocalisation following optogenetic activation, validating the use of equation (1).

When we examined the temporal evolution of myosin and tension more closely, we observed that, following the peak in activation, the signal for MRLC-iRFP followed a different trend than CRY2-p115RhoGEF and tension (**Fig 5d-f**). Indeed, after reaching their peak, CRY2-p115RhoGEF and tension decayed over ∼400s, whereas myosin decayed significantly more slowly. Intriguingly, such an effect was not observed in interphase cells (**Fig 2a-c**) and may indicate that myosin localisation is not perfectly correlated with contractility in mitosis, perhaps because the dense metaphase cortex prevents penetration of mini-filaments newly recruited from the cytoplasm into the cortex^32^. How the kinetics of tension generation is controlled by regulation of myosin activity and penetration of mini-filaments within the cortex will be interesting to explore.

Overall, the couples *(κ*_*myosin*_, *τ*_*myosin*_) and *(κ*_*tension*_, *τ*_*tension*_) quantify how recruitment of p115RhoGEF to the membrane leads to myosin recruitment and cortical tension change.

### Local changes in cell shape can be predicted based on relationship between signalling and tension

Overall, our experiments imply that it may be possible to deduce what mechanical changes take place in the cell simply by observing the spatiotemporal pattern of RhoGEFs at the membrane. As a test of this idea, we investigated if we could predict the cellular shape changes induced by localised recruitment of CRY2-p115RhoGEF to the membrane of rounded mitotic cells blocked in metaphase. In rounded cells, a localised increase in RhoA activity should lead to a local increase in cortical tension and local flattening. Such shape changes are observed for example in anaphase when myosin accumulation at the cell equator leads to a localised flattening prior to the onset of furrowing ^33^.

In our experiments, we exposed a small region of the membrane of a mitotic cell expressing CRY2-p115RhoGEF to 300ms pulses of blue light repeated at 2-minute intervals. This resulted in a steady level of recruitment of CRY2-p115RhoGEF to the membrane in the region of activation and subsequently gave rise to localised recruitment of myosin (3/4 cells, **Fig 6a**, 1/4 cell (not shown) showed limited myosin recruitment following activation and no deformation). Over time, the cells changed shape and became visibly flattened in the region of stimulation (3/4 cells, **Fig 6a**,**b**), as expected if tension was locally higher.

**Figure 6.**
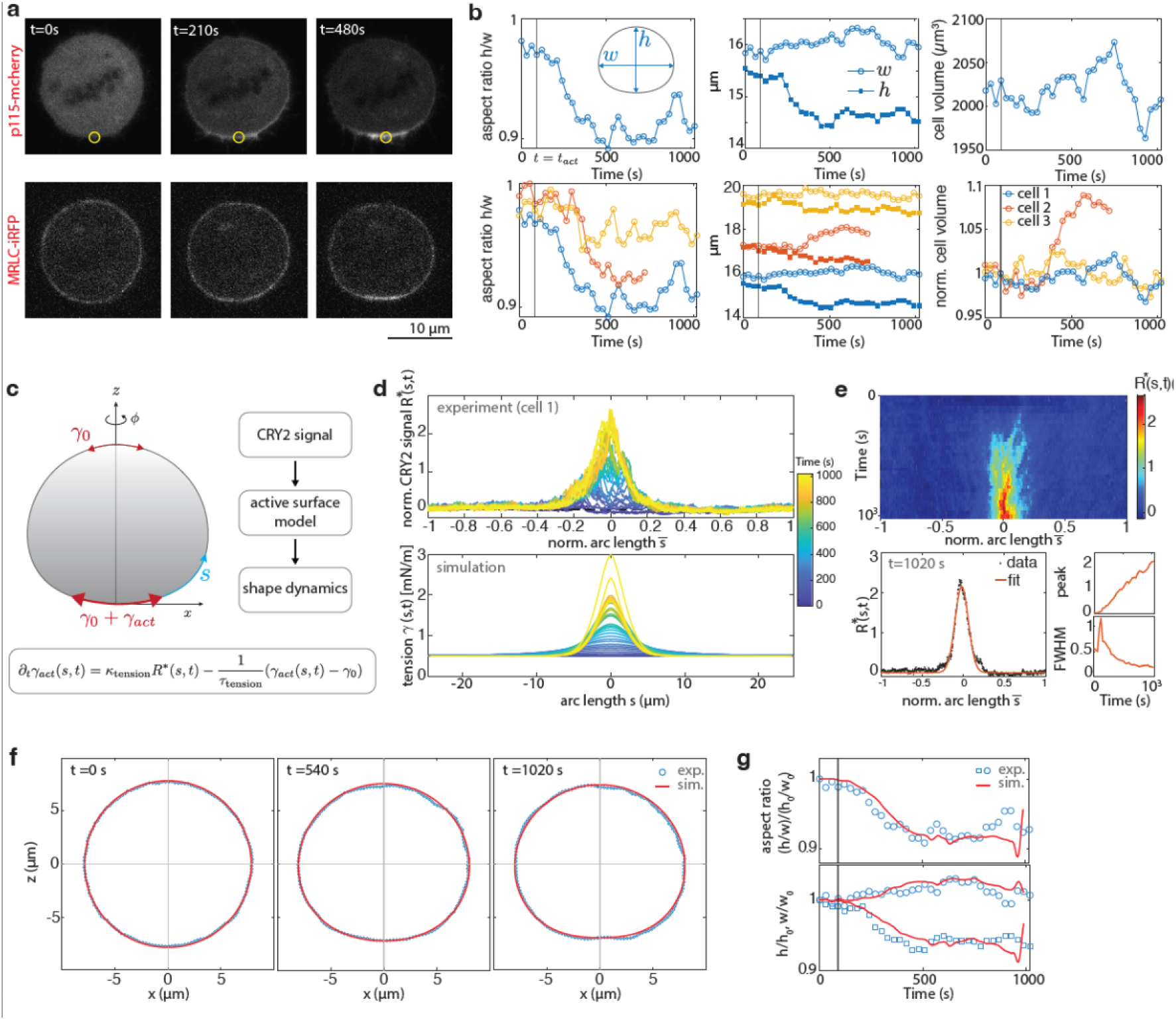
Coarse-grained characterisation of the mechanical output of signalling allows prediction of cell shape changes. **a.** Example of cortical activation near the cell south pole (yellow circle). The top row shows the p115-mCherry signal and the bottom row the MRLC-iRFP signal. **b**. Temporal evolution of cell aspect ratio *h*/*w*, cell height and width *h* and *w*, and cell volume, following localised activation at *t*=*t*_*act*_. Top: one example cell (cell 1 in the bottom graphs), bottom: overlay of data for 3 activated cells. **c**. Schematic of active surface model of cell shape dynamics. The profile of CRY2 at the membrane is converted into a tension profile using the input output relationship from equation 1 (shown below the diagram) and *τ*_*tension*_ from Fig 5i. This then serves as input into an active surface model to predict shape dynamics. **d**. Top: Experimental measurement of profile of CRY2* as a function of the normalised arc length 0 along the cell contour, color-coded according to time. Bottom: tension profile predicted as a response to CRY2* profiles fitted as shown in panel e with *τ*_*tension*_ =30s and *κ*_*tension*_ = 0.025 mN.m^-1^.s^-1^ (see Supplementary Information). **e**. Kymograph of experimentally measured CRY2 profile (same data as in panel d), example of von Mises fit at one time point and plot of fitted peak values (top) and full-width-at-half-maximum (FWHM, bottom) over time. **f, g**. Overlay of experimental result (blue circles) and simulation dynamics (red line) of the cell shape (f) and dynamics of normalised aspect ratio, height and width as a function of time (g), for one representative example (cell 1).

Next, we sought to predict the cellular shape change from knowledge of the spatiotemporal pattern of signalling, based on the delayed response of tension found in our experiments of fully activated cells (**Fig. 2, Fig. 5**). The spatiotemporal profile of CRY2-p115RhoGEF recruitment to the membrane upon localised activation was used as an input into a mechanical simulation of the cell surface^34^ (**Fig 6c, Supplementary Fig S2**, and **Supplementary Information**). With this approach^34-36^, the cell surface is considered as an active viscous gel parametrised with an active tension *γ* and a viscosity *η*. The active tension was taken as the sum of a uniform reference tension *γ* _0_, and a spatially varying tension *γ* _*act*_ (*s,t*) · *γ* _*act*_ (*s,t*) locally evolves in time as a response to activated CRY2*, according to Equation (1), with a delay *τ*_*tension*_ and a rescaling factor *κ*_*tension*_ (**Fig 2, Fig 5, Fig 6c**). In addition, the cell volume of the simulated cell was taken from experimental measurements of inferred cell volume, assuming axisymmetry around an axis passing through the optogenetically activated domain (**Fig 6b**). For each experiment, the simulation predicts the temporal evolution of cell shape in response to the temporally varying spatial profile of CRY2-p115RhoGEF fluorescence. The reference tension was set to the average tension measured in mitotic cells, *γ*_0_ = 0.5 mN.m^-1^ (**Fig 5b**), we imposed *κ*_*tension*_=30s, as measured from full cell activation experiments (**Fig 5**), and a cortical viscosity of *η*= 3 ×10^4^ Pa.µm.s (**Supplementary Fig S3, Supplementary Tables S1-3, Supplementary results**). As the value of R* in local activation experiments was approximately an order of magnitude larger than in global activations (**Fig 6d, Fig 5d, Supplementary Table S2**), we decided to adjust the rescaling factor *κ*_*tension*_ for each cell. We found that the predicted dynamics of cell shape agreed well with experimentally measured cell deformations (**Fig 6 f**,**g, Supplementary Information**). The rescaling factors *κ*_*tension*_ were however ∼7 times smaller than measured from full cell activation, indicating that CRY2 is less efficient at triggering tension increase in local activation experiment. Since the localized cell activation leads to local values of CRY2 much larger than in full activation experiments (compare **Fig 5d** and **6d**), we surmise that this difference arises from saturation at high CRY2 concentration of the linear relationship between tension increase and membranous CRY2 increase observed in **Fig 5j**.

In summary, characterising the mechanical response of the cell to a pulse in signalling allows prediction of the spatiotemporal evolution of cell shape in response to sustained localised changes in RhoGEF signalling.

## Discussion

### A quantitative framework linking RhoGEF signalling to cortical mechanics and shape

RhoGTPase signalling controls actin cortex mechanics during cell and tissue morphogenesis. Here, we characterised the relationship between the amount of membranous RhoGEF and the downstream molecular and mechanical changes using optogenetics. We developed a predictive model of the temporal evolution of RhoGEF membrane localisation in response to a pulse of light. We showed that myosin recruitment to the cortex and increase in cortical tension occur with a delay from the time of RhoGEF localisation to the membrane. A second predictive model uses the temporal evolution of membranous RhoGEF to predict the temporal evolution of downstream outputs of signalling, such as myosin recruitment to the cortex and cortical tension. We validated this model and showed that cortical myosin and tension both increase linearly with the amount of signalling proteins at the membrane. Using the characterisation of the mechanical changes induced by RhoGEF signalling, we were able to predict the shape changes induced by localised recruitment of signalling proteins, directly linking gradients in signalling to shape change. In summary, we provide a theoretical and experimental approach to link signalling to mechanical changes and shape changes that has broad applicability.

### Mechanical changes scale linearly with the amount of RhoGEF signalling

After activation, RhoGEFs localise to specific subcellular compartments to modulate RhoGTPase activity locally ^30^. We characterised myosin recruitment and the mechanical response to relocalisation of the DH-PH domain of RhoA-specific GEFs to the cell membrane. In response to a single pulse of optogenetic actuation, the evolution of myosin recruitment and cortical tension followed similar temporal patterns to the localisation of the RhoGEF domain to the membrane except with a delay. This suggested that, in our experimental conditions, all steps in the cascade linking RhoGEF to myosin and tension were linear. By systematically varying illumination, we showed that myosin recruitment to the cortex and cortical tension increase scale linearly with the amount of RhoGEF recruited to the membrane. For any output *O*(*t*) downstream of signalling, the couple (*κ*_o_, *τ*_o_) quantify the response of the cell.

One limitation of our current work is that we only examined rounded cells. This was dictated by the need to simultaneously quantify relocalisation of signalling to the membrane and measure cortical tension. However, our approach is generally applicable to any situation in which the temporal relocalisation of an optogenetic actuator can be quantified together with an output (mechanical or biological), for example the traction force exerted by optogenetically-controlled epithelial cells ^17,18^.

Our experiments examined the mechanical response of the cell to a single pulse of light. Indeed, for a linear system, knowledge of the system response to an activation pulse enables the prediction of response to arbitrarily complex patterns of activation via convolution. However, future work will be necessary to understand the response to more prolonged signals and understand the conditions in which the cell behaves as a linear system. Such experiments may reveal saturation of signalling or lead to the activation of additional effectors. For example, when we locally recruited RhoGEFs to the membrane of mitotic cells, the membranous concentration of CRY2* was one order of magnitude larger than in global activation experiments but, in contrast, the rescaling factor *κ*_*tension*_ was ∼7 fold smaller. This may be because the RhoGEFs compete locally for a limited substrate, RhoA-GDP. In our experiments, we could not detect any changes in F-actin in response to single pulses of optogenetic activation, whereas experiments applying constant illumination over several minutes reveal extensive actin remodelling ^13^.

Intriguingly, our experiments also revealed that cortical myosin enrichment and tension took longer to return to their baseline in metaphase than in interphase. Our optogenetic system only allows control over activation of signalling and, therefore, studying the return to initial values may allow investigation of the molecular mechanisms controlling tensional homeostasis. One implication of our work is that it may be possible to deduce what cytoskeletal and mechanical changes take place in the cell simply by observing the localisation and enrichment of signalling proteins controlling RhoA activity, such as RhoGEFs or RhoGAPs. In support of this idea, we were able to predict the cellular shape changes induced by sustained localised RhoGEF signalling from characterisation of the cellular mechanical response to single pulses in signalling. However, further work will be necessary to fully explore this.

### Coarse graining the mechanical response to signalling

Signalling cascades are complex and involve a large number of proteins, whose abundance and activation state are not known, that interact with partners, with reaction constants that are often poorly characterised. Therefore, predicting the outcome of changes in signalling is complex and necessitates coarse-graining approaches.

Our experiments showed that cytoskeletal and mechanical changes could be deduced from the temporal evolution of the localisation of RhoGEF to the membrane through introduction of a delay and a rescaling. We showed that myosin recruitment and tension increase scale linearly with the amount of signalling proteins at the membrane. This suggests that, in our experimental conditions, downstream signalling can be deduced from spatiotemporal changes in an input signal – the localisation of the RhoGEF to the membrane. The effect of signalling can be coarse grained by a pair of parameters (*κ*_*o*_, *τ*_*o*_), specific to each output.

As we showed linear scaling between input and output, it may be possible to coarse grain the output of complex signalling pathways in response to a change in signalling by using the concept of transfer functions. Transfer functions *T*(*x,t*) originate from signal processing and they describe how a system’s output relates to an input signal. In our context, the temporal evolution of RhoGEF localisation at the membrane constitutes the input signal. RhoGTPase activation, remodelling of the cortex and myosin activation downstream of this input participate in setting the transfer function. The resulting changes in cortical mechanics constitute the output signal. In our experiments, we stimulate signalling with a pulse of light, akin to a Dirac function. As a consequence, the output that we measure for change in cortical tension represents an approximation of the transfer function linking the light input to the mechanical output of signalling. In principle, convolution of this transfer function with a more complex pattern of light stimulation should allow prediction of the mechanical output of the system in response to an arbitrary input signal, as long as no part of the signalling cascade saturates. In future, it will be interesting to explore the generality and limitations of such an approach, in particular identifying the conditions that lead to saturation. Combining optogenetic actuation with imaging of live reporters of protein activity should allow dissection of the different steps in a signalling cascade and identification of which protein components saturate.

Further, through combining the measured delay dynamics of tension with a mechanical model, we have a tool to predict cellular shape dynamics from localised changes in signalling induced optogenetically. Using spatio-temporally patterned optogenetic stimuli would thus allow us to programme specific cell shape deformations. Investigating more pronounced shape changes and their reversibility properties in the future will refine this tool for controllable morphogenesis in synthetic biology applications^37^.

### Towards a biophysical signature of RhoGEFs

RhoGEFs exhibit significant diversity, with many acting on one or multiple RhoGTPases ^30,38^. In particular, previous work has shown that a same RhoGEF can induce distinct responses dependent on the RhoGTPase it interacts^31^. However, quantifying the impact of RhoGEFs on mechanics is challenging, even when they act on the same RhoGTPase. A potential approach to classifying RhoGEFs is to characterise their impact on cell mechanics. Our experiments show that two scalar parameters, the amplitude *κ* and the delay *τ* defined in equation (1), may provide a first signature of the regulatory effect of RhoGEFs on mechanics. Our experiments revealed that two closely related GEFs p115RhoGEF and PDZRhoGEF had similar signatures but differed from Ect2 that is more distantly related^6^. A more systematic use of this approach may help to understand why such a wide variety of RhoGEFs has evolved, potentially allowing us to correlate mechanical signature with biological function. Furthermore, new high throughput approaches for characterising mechanics based on flow cytometry ^39,40^ will enable more rapid characterisation of mechanical signatures of modulators of RhoGTPase activity.

## Materials and methods

### Cloning

The plasmids pCRY2PHR-mCherryN1 and pCIBN(deltaNLS)-pmGFP were acquired from Addgene (plasmid #26866, and plasmid #26867) ^23^. The plasmid for ARHGEF1-DH-PH-CRY2PHR-mCherry is described in ^17,24^. The DH-PH domains of ARHGEF11 (PDZ-RhoGEF) and Ect2 were identified using Uniprot. We extended the sequence of interest to retain 8 extra amino acids at either extremity of this catalytic domain following a previously used approach ^12^. This domain was inserted into the CRY2-mCherry plasmid using restriction digest. The obtained plasmids ARHGEF11-DH-PH-CRY2PHR-mCherry and Ect2-DH-PH-CRY2PHR-mCherry were then inserted into a lentiviral vector using the Gateway technology first into a *p-DONR*-221 vector and finally in a *p-DEST*-Neomycin vector (Thermo Fisher). CIBN(deltaNLS)-pmGFP was inserted in a retroviral vector pRetroQAcGFP-N1 (Clontech). MRLC-iRFP and LifeACT-iRFP plasmids were a kind gift from Leo Valon (Institut Pasteur, Paris, France). Anillin-AHD-PH-iRFP (RhoA biosensor) is described in ^24^. All clones were verified by sequencing (SourceBioscience). All plasmids contain the generic CMV promoter from the backbone pmCherry-N1 (Clontech Laboratories). All plasmids are deposited in Addgene.

### Cell culture and generation of cell lines

MDCKII cells were cultured at 37°C with 5%CO_2_ in high-glucose (4.5g/L) DMEM (Thermo Fisher) supplemented with 10% fetal bovine serum (Sigma), 1% penicillin-streptomycin (Thermo Fisher) and 25mM Hepes buffer (Gibco). Where appropriate, the medium was supplemented with selection antibiotics: G418 1mg/mL (Sigma), puromycin 1µg/mL (Calbiochem) and HygromycinB 300µg/mL (Invivogen). A stable cell line expressing CIBN-GFP-CAAX was made by retroviral transduction into MDCKII WT cells. Cell lines stably expressing CRY2-mCherry, ARHGEF1-CRY2-mCherry, ARHGEF11-CRY2-mCherry, Ect2-CRY2-mCherry, MRLC-iRFP, LifeACT-iRFP or iRFP-RhoA biosensor were made by lentiviral transductions. All cell lines were selected with appropriate antibiotics and sorted by flow cytometry before use.

Cells were routinely tested for the presence of mycoplasma using the mycoALERT kit (Lonza). None of the cell lines in this study were found in the database of commonly misidentified cell lines maintained by ICLAC and NCBI Biosample.

All imaging was done in Leibovitz L-15 medium (Gibco) supplemented with 10% fetal calf serum or in phenol-red-free DMEM (Gibco) supplemented with 10% fetal bovine serum, L-Glutamine and penicillin/streptomycin.

### Flow cytometry

Flow Cytometry was carried out on a FACSAria III. Cells were grown to full confluence in a T75 flask (∼10^7^ cells). Cells were resuspended in 1.5 mL of Gey’s balanced salt solution (Sigma) and passed through a cell strainer (40µm mesh size). Cells were sorted using the appropriate laser for fluorescence activation of EGFP, mCherry, or iRFP. Only the cells with the highest 5% fluorescence were collected. After sorting, the cells were resuspended in normal growth medium and amplified for use in experiments.

### Sample preparation for tension measurement

Measurement of cortical tension requires rounded cells seeded at low density onto a substrate. The standard protocol to measure the tension of interphase cells involves detaching cells and seeding them at low density onto petri dishes just before the experiment. However, we observed significant loss of CRY2-mCherry fluorescence with this approach, preventing our experiments.

Therefore for interphase cells, we seeded single cells onto fibronectin coated circular micropatterns 10μm in diameter 15–20 hours prior to the experiment. This approach avoided CRY2 Each step of the process of creating the micropatterns is shown in **Supplementary Fig S4a**. Briefly, glass bottom petri dishes (35 mm diameter, World Precision Instruments) were coated with Trichloro(1H,1H,2H,2H-perfluorooctylsilane) by vapor deposition at 0.3 mbar for 30min. After silanisation, a 0.2% Pluronic F-127 solution in water was applied for 30 min to block protein adsorption. A PDMS stamp with 10μm circular cross-section pillars was incubated with a fibronectin solution (10μg/mL), dried, and gently pressed onto the Pluronic coated glass to generate circular fibronectin patterns. The surface was rinsed with PBS, leaving the substrate ready for cell culture. Cells were seeded onto the micropatterns and incubated for 15–30 minutes. Non-adherent cells were removed by gently rinsing with cell medium.

For experiments on mitotic cells, ∼20,000 cells were plated on 35mm glass bottom dishes (World Precision Instruments) ∼36 hours prior to imaging. On the day of the experiment, they were treated with Nocodazole (30 ng/mL) for 2–3 hours, washed twice with drug-free medium to remove residual Nocodazole, and incubated with MG132 for 10 minutes to arrest them in metaphase.

### Atomic Force Microscopy

#### Cantilever calibration and bead attachment

Tension measurements were performed using a tipless silicon cantilever (ARROW-TL1Au-50-NanoWorld) with a nominal spring constant of 0.03N/m mounted on a JPK CellHesion module (JPK instruments) on an IX81 inverted confocal microscope Olympus-FV1000 (Olympus). Imaging was done with a 63x oil immersion objective (Olympus, N.A.=1.42).

A 150μm radius bead was attached to the cantilevers prior to the measurements to achieve a pseudo-parallel plate compression setup (**Supplementary figure S5a**). Each step of the process of bead attachment is shown in **Supplementary Fig S4b**. Prior to attachment, the cantilever’s spring constant was determined using the thermal fluctuation method, with an average value of 0.046±0.023N/m. Next, a 150μm latex bead was attached to the cantilever using NOA UV glue (Edmund Optics). The underside of the cantilever was briefly brought into contact with a small glue patch spread on a glass slide. Then the cantilever was positioned above an isolated bead and brought into contact with it. The bead and cantilever were exposed to UV light to cure the glue, securely bonding the bead to the cantilever. Due to the large radius of the bead relative to the cell size, the bead surface curvature was minimal over the contact area, approximating a flat plate (**Fig 1e**). For example, for a compression radius l = 5μm (half of the typical cell diameter) with a bead of radius R=150 μm, the sagitta *s* of the spherical cap was calculated as: 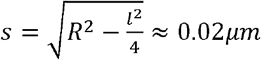 demonstrating near-flatness at the contact interface.

In experiments, a first force-distance curve was acquired on the glass surface close to the cell of interest to calibrate the sensitivity of the cantilever and the position of the substrate surface relative to the retracted cantilever (h_cantilever_). A second, force-distance curve was acquired on top of the cell allowing determination of the cell radius R_c_.

### Principle and theory of cortical tension measurement

To measure cortical tension and its response to optogenetically triggered GEF localisation to the membrane, we compressed single rounded cells with a large bead attached to an AFM cantilever. The cell’s response to compression exhibits shell-like behavior ^41^, characterized by an area compressibility modulus (K_A_) and a pre-stress term (T_0_), which describe the mechanical properties of the cortical shell. Before compression, the cell adopts a spherical shape with radius Rc (**Supplementary Fig S5a, left**). Upon compression with the bead, the cell shape becomes a spherical cap (**Supplementary Fig S5a** right), resulting in an increase in surface area. The elastic nature of the cortex at subsecond timescales resists this deformation, leading to a transient increase in internal stress and a peak in the force measured by the AFM cantilever. An example of AFM force curve is shown (**Supplementary Fig S5b**). Over the following 30s, the viscoelastic properties of the cortex that arise due to biochemical turnover of cortical proteins dissipate this stress, resulting in a force relaxation phase that eventually stabilises to a plateau, representing a steady-state ^26^. This plateau corresponds to the preactivation cortical tension (T_0_), which reflects the baseline mechanical tension in the cortex.

The force on the AFM cantilever during relaxation can be written as ^26,41^:

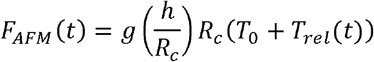

where Rc is the cell radius before compression (**Supplementary Fig S5a**, left), T_0_ the cortical tension, 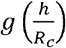 a generic function that only depends on the compression height h and Rc . It is expressed as:

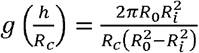

with R_0_ the equatorial radius and R_i_ the contact radius (**Supplementary Fig S5a**, right). Since R_i_ is challenging to measure experimentally, g is calculated numerically from volume conservation and force balance equations and approximated using a third-order polynomial fit: 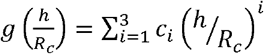 . For more detail, see the theory section in the supplementary materials from ref ^41^.

T_rel_ is the time-dependent tension describing the stress relaxation and T_rel_ converges to 0, when the relaxation reaches the plateau. The original article^41^ incorporates a time-dependent power-law beahaviour to model cortical relaxation after compression:

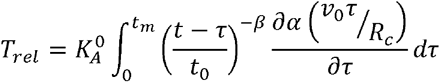

With t_m_ the time marking the beginning of relaxation and *ν*_0_ the cantilever’s approach velocity. An example of a fit of F_AFM_(t) is shown in **Supplementary Fig S5b**. For t_0_ (a time just before the optogenetic activation time t_act_), the stress due to the compression is completely dissipated and the force simplifies to:

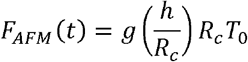

Optogenetic activation at t_act_ induces an increase in cortical tension Δ*T* that results in an increase in the force detected by AFM. After activation, the AFM force is given by:

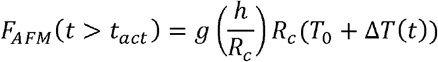

We obtain the following equation s to calculate the pre-activation cortical tension and the change on cortical tension after optogenetic activation:

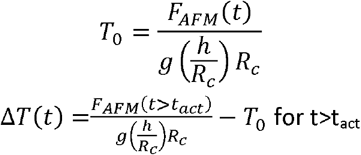

An example of tension variation after optogenetic activation, Δ*T* (*t*), is shown in **Supplementary Fig S5c** and the normalised tension 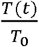 is shown in **Supplementary Fig S5d** for a basal tension of T_0_=0.47 mN/m.

### Cortical tension measurement during optogenetic activation

The approach speed of the cantilever was set at 10µm/s. We chose a set-point force of 15nN, which produced an average cell compression of 2.5 - 4.7 μm, which is 24-35% of the cell height. During the constant height compression, the force acting on the cantilever was recorded. After initial force relaxation of ∼40sec, the resulting force value was used to extract surface tension pre-activation (T_0_). At about ∼80sec after the onset of compression, optogenetic activation was carried out on Olympus FV-1000, using the 488-nm light at 10% laser power (2.3mW).

### Drug treatments

Drug treatments were performed as follows. To inhibit myosin contractility, s-nitro-blebbistatin (Cayman Chemicals) was added at 20µM concentration 10mins before experiments. We used this more stable form of blebbistatin as it has been reported that addition of a nitro group stabilizes the molecule thus preventing its degradation when exposed to 473-nm light ^42^.

### Optical Microscopy and optogenetic activation

Imaging experiments were performed on an Olympus IX83 inverted microscope equipped with a scanning laser confocal head (Olympus FV-1200). Combined mechanical measurements and optogenetic activation experiments were performed on an Olympus IX81 inverted microscope equipped with an Olympus FV-1000 scanning laser confocal head. All images were acquired with a 60x oil immersion objective (NA 1.35, Olympus). For brightfield imaging, a far-red filter was inserted into the illumination path to avoid spurious activation of CRY2. Images were acquired at the equatorial plane of rounded cells.

Optogenetic activation was carried out with the 473nm laser in a circular region at the centre of the cell with a radius r_act_∼200nm illuminated with a spiral movement. During experiments, we acquired images at 1-2s interval for durations of 500s. Imaging of CRY2-mCherry was performed using a 559-nm laser and imaging of MRLC-iRFP, LifeACT-iRFP,and the iRFP-RhoA biosensor was done using a 633-nm laser. In imaging experiments on the FV1200, a GaAsP-high sensitivity detector unit was used to image mCherry and iRFP signals and reduce laser exposure times. At the end of each experiment, we also acquired an image of the CIBN-GFP-CAAX signal in each cell.

### Image processing

A detailed description of image processing is given in supplementary methods.

### RNA extraction and Quantitative Real-Time PCR

Total RNA from the relevant MDCK cell lines was extracted using the RNeasy Mini Kit (Qiagen, Hilden, Germany) and reverse transcribed using the High Capacity cDNA Reverse Transcription Kit (Applied Biosystems, Carlsbad, CA) following manufacturer protocols. The gene expression level for endogenous controls *GAPDH* (Hs00266705) and *HPRT1* (Hs00357333) was determined using pre-validated Taqman Gene Expression Assays (Applied Biosystems), and gene expression level for genes of interest (GFP, mCherry, RHOA, MYH9, MYH10) was determined using assays designed with the Universal Probe Library (UPL) from Roche (www.universalprobelibrary.com) according to manufacturer’s instructions. Sequences of primers used are available upon request. Gene expression was normalized to *HRPT1* and *GAPDH* expression.

### Curve fitting, statistical analysis, and reproducibility

All experiments were repeated on at least two days with similar results. The exact number of repeats for each experiment is indicated in the appropriate figure legend. Wilcoxon rank sum test. Outliers were identified using the median absolute deviation (MAD) method with MATLAB. Outliers were defined as values that exceeded three scaled MADs from the median. The scaled MAD is calculated as a constant multiplied by the median of the absolute deviations from the median of the data. Detected outliers were excluded from the statistical analysis.

## Code Availability

The code utilized to analyse data in this study is available from the authors upon request.

## Data and reagent Availability

All data supporting the conclusions here are deposited in the UCL data repository with a unique doi (https:\\rdr.ucl.ac.uk).

## Acknowledgements

We thank Mathieu Coppey (Institut Curie, Paris, France) for advice on optogenetics and sharing reagents, Ayad Eddaoudi (UCL, Institute for Child Health) for flow cytometry analysis, and members of the Charras and Salbreux labs for discussions. PB and DK were supported by a CRUK multidisciplinary award (C55977/A23342) to GC and GS. MK was supported by a SNSF early post-doc fellowship P2LAP3_164919. MK and EF were supported by a European Research Council consolidator grant (CoG-647186) to GC. GC was supported by grant BB/W011123/1 from the Biotechnology and Biological Sciences Research Council. GS was supported by the Francis Crick Institute which receives its core funding from Cancer Research UK (FC001317), the UK Medical Research Council (FC001317) and the Wellcome Trust (FC001317).

## Author contributions

P.B., G.S. and G.C. conceived the project and wrote the manuscript. P.B. performed most of the experiments and analysis. P.B. designed the plugin for segmentation analysis. M.K. generated cell lines and performed some experiments. E.F. and L.V. helped in flow cytometry and generation of plasmids. P.P.R. and G.L. performed the qRT-PCR experiments. D.K. and G.S. designed the computational model for cellular shape prediction. D.K. performed cell shape prediction with the computational model. G.S. contributed to model design and data analysis. G.C. oversaw the entire project and writing. All authors discussed the results and manuscript.

