## Supplementary material for "Control of cellular cortical tension and shape by RhoGTPase signalling"

December 12, 2025

### 1 Image processing

#### 1.1 Determination of radial profiles of CRY2 fluorescence intensity

Image segmentation and contour detection were performed using procedures implemented in Matlab. At each time point and for each image of the movie, the cell contour was detected (Fig S1a-1) using active contours based on the level set method [3, 14]. When the nucleus was visible within the plane of imaging, the same method was used on an average image of all of the frames of the movie[1] to obtain the contour of the nucleus (Fig S1a-2) to exclude its contribution from fluorescence measurements.

Next, we used the pixels of the contour to calculate the average fluorescence intensity along the cell periphery. To obtain the average radial profile of the fluorescence intensity of CRY2, we used the cell contour to generate a mask, denoted $B_0$, such that a pixel has a value of 1 if it is inside the contour and 0 outside (Fig S1a-3). Then, this binary image was eroded to obtain $B_{-1}$ (Fig S1a-4) corresponding to the pixels located inside $B_0$. By taking the difference between the binary image $B_0$ and the binary image $B_{-1}$, we were able to obtain the pixels located in the area between these two masks, $C_0$, (Fig S1a-5), which corresponds to the cell boundary. Finally, we computed the average fluorescence intensity of pixels in $C_0$ and $B_{-1}$ to obtain the cortical fluorescence and the cytoplasmic fluorescence, respectively.

To determine the position of the cell centre, we iterated eroding operations $B_{-i} \to B_{-i-1}$. This yielded concentric rings of decreasing radius, $C_{-i}$. Iterations were stopped when the binary image was fully eroded, leaving a single pixel which we identified as the cell centre. A reverse operation was performed to define iteration steps $B_i \to B_{i+1}$ [2], giving a set of contours outside of the cell, $C_i$. This operation was stopped when the edge of the image was reached.

From these operations, we obtain a family of contours $C_i$ at increasing distance from the cell centre. We then calculated the average intensity within each contour to obtain the radial profile of CRY2 fluorescence intensity in the cell $I(x,t)$ at each time $t$ as a function of the distance $x$ from the cell centre before and after the optogenetic activation. Fig S1b shows the radial fluorescence profile of CRY2 before optogenetic activation and Fig S1c the profile 45s after activation.

We now describe procedures to normalise these fluorescence intensity profiles and correct for bleaching. First, the average intensity of the background was subtracted from the radial profile : $\Delta I(x,t) = I(x,t) - I_{background}(t)$ where $I_{background}(t) = \langle I(x,t) \rangle_{x_{out}}$ is the average fluorescence intensity for all radii outside the cell at a given time $t$. Then, the effect of photobleaching[3] was corrected for by dividing each pixel within the cell by the average fluorescence intensity of the whole cell $\langle \Delta I_{cell} \rangle (t)$ (i.e the average intensity of the pixels inside the contour, for which the background was also subtracted). Indeed, we reasoned that, over the timescale of our experiment, the total number of CRY2-mCherry in the cell remains constant. We note however that $\langle \Delta I_{cell} \rangle (t)$ is only approximately proportional to the total cell fluorescence, as we image only one confocal plane. Fig S1d displays the normalised radial profile $i(x,t)$ for each time $t$ :

$$i(x,t) = \frac{I(x,t) - I_{background}(t)}{\langle \Delta I_{cell} \rangle (t)} \tag{1}$$

In Fig S1d, the black curves show the radial profiles of normalised intensity before optogenetic activation in the cell of interest. Before activation, fluorescence is approximately uniform as a function of distance within the cell, as expected for a cytoplasmic fluorophore (Fig S1b). The grey curves show the normalised radial profile after the activation and exhibit a sharp peak at the cell boundary, as expected for a fluorophore that relocalises to the cell membrane (Fig S1c).

[1] We resorted to the average as obtaining the nuclear contour (inset Fig S1c) for each frame was challenging, because of the decrease in cytoplasmic fluorescence intensity induced by optogenetic relocalisation of CRY2 to the plasma membrane.

[2] In this case, we used the dilate function and the difference calculation is between the binary images $B_{i+1}$ and $B_i$

[3] For a homogeneous illumination, bleaching is a function of time $t$ and laser power $P_{laser}$ : $f_{bleaching}(t, P_{laser})$. A practical way to remove the effect of photobleaching is to divide the intensity by a variable that depends on photobleaching in the same manner as the variable of interest.

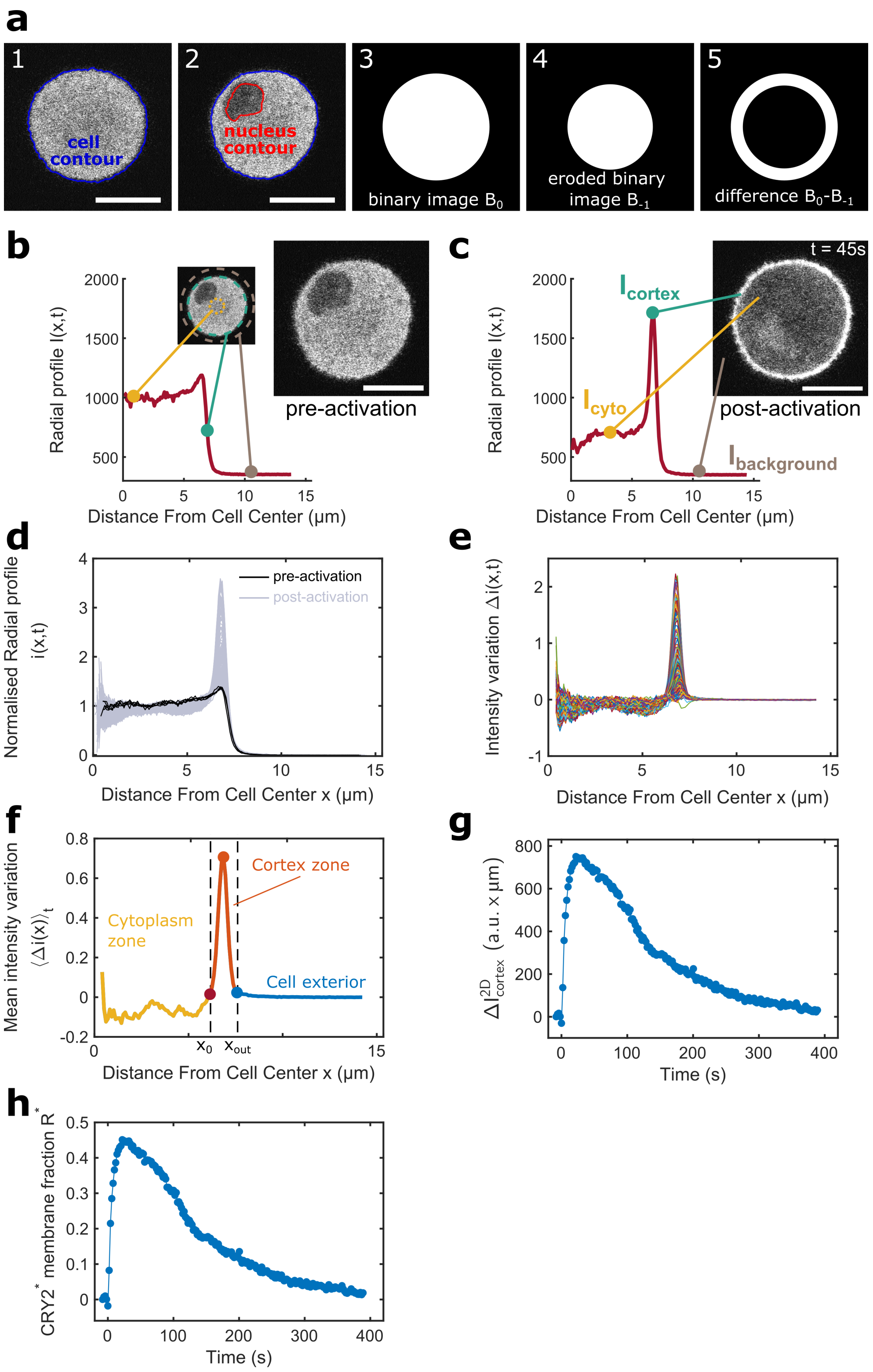


Supplementary Figure 1: **(a)** Steps of the image processing procedure. All examples in the figure are based on the same representative cell to illustrate the procedure. **(a1-2)** Image of the cell with its detected contour and, when visible, the nuclear contour (a2). The contours were obtained using active contour detection based on the level set method. **(a3)** Example of a binary image $B_0$, where pixels inside the cell contour are set to 1, and those outside are set to 0. **(a4)** The eroded binary image $B_{-1}$, which represents the inner shape of $B_0$. **(a5)** The difference between $B_0$ and $B_{-1}$, highlighting the pixels at the cell boundary.

Supplementary Figure 1 (continued): **(b-g)** Steps of the quantitative analysis of fluorescence intensity before and after optogenetic activation. **(b, c)** Images of CRY2-mCherry localisation and radial profile of fluorescence intensity $I(x,t)$ as a function of the distance from the cell centre before (b) and 45s after (c) optogenetic activation. The white scale bars represent 10 $\mu$m. Key regions of the radial fluorescence intensity curves are shown. **(d)** Normalised radial profile $i(x,t)$, obtained after background subtraction and correction for photobleaching before (black) and after (grey) optogenetic activation **(e)** Temporal intensity variation $\Delta i(x,t)$ at each radial position, calculated as the difference between the normalized intensity before and after optogenetic activation. **(f)** Mean intensity variation during the recruitment phase, used to define the cytoplasmic (yellow) and cortical regions (orange), as well as the cell exterior (blue). The cortical region is defined as the zone where fluorescence intensity increases upon activation, while the cytoplasmic region corresponds to the area where fluorescence decreases. **(g)** Temporal evolution $\Delta I^{2D}_{\text{cortex}}(t)$, considered to be proportional to membranous $[\text{CRY2}^* - \text{CIBN}]$ (blue). **(h)** Temporal evolution of proportion of activated CRY2 at the membrane, $R^*$.

To measure the effect of optogenetic activation, we calculated the temporal change of normalised intensity $\Delta i(x,t)$ between the normalised intensity before and after optogenetic activation (Fig S1e), at each distance $x$:

$$\Delta i(x,t) = i(x,t) - i_{preact}(x) \tag{2}$$

where $i_{preact}(x) = \langle i(x,t) \rangle_{t<t_{act}}$ is the temporal average of the radial profiles before the activation time $t_{act}$ [4].

#### 1.2 Averaging within cortical and cytoplasmic regions

After activation by blue light, the optogenetic protein CRY2 undergoes a conformational change and, when it encounters the cell membrane after diffusing through the cytoplasm, it can bind the membrane localised CIBN-CAAX. As a result, CRY2-mcherry is depleted from the cytoplasm and enriched at the cortex. Therefore, we expect the fluorescence signal to increase in the cortex and decrease in the cytoplasm. We used two different methods to analyze variations in the fluorescence signal due to photoactivation.

To provide a first estimation of the amplitude and kinetics of recruitment of CRY2 to the cortex, we calculated the average intensity profile away from the center of the cell $I(x,t)$, with $x$ the distance from the center. Then, we defined the cortical fluorescence intensity as the maximum value of the shifted radial profile $\Delta I_{cortex}(t) = I(x_{max},t) - I_{background}(t)$ with $x_{max}$ the radial position at which intensity is maximal at time t (Fig S1c). The cytoplasmic intensity was defined as the average along $x$ of the radial profile for $x$ contained within the cell (Fig S1c). For this, we defined the position of the outer boundary of the cytoplasm as $x_{cyto} = x_{max} - 10$ pixels and started averaging 5 pixels away from the centre of the cell to minimise the contribution of noise due to averaging over a small number of pixels in the inner rings. This yields an averaged value for fluorescence intensity in the cytoplasm : $\Delta I_{cyto}(t) = \langle I(x,t) \rangle_{x\in[5,x_{cyto}]} - I_{background}(t)$. In Figs 2a-b, we used the ratio $\frac{\Delta I_{cortex}}{\Delta I_{cyto}}$ normalised to its value at $t = t_{act}$ to quantify the temporal evolution of CRY2 fluorescence at the membrane during the optogenetic experiments.

Second, we devised an alternative method to more precisely define a cytoplasmic compartment (characterized by a loss of fluorescence intensity following activation) and a cortical, surface compartment (characterized by recruitment of the fluorescent probe). To distinguish cortex from cytoplasm, we used the normalized fluorescence intensity variation $\Delta i(x,t)$ during the recruitment phase ($t <$ 50s after activation). We reasoned that a point at position $x_{cyto}$ is in the cytoplasm if $\Delta i(x_{cyto},t) \leq 0$, and a point at position $x_{cortex}$ is in the cortex if $\Delta i(x_{cortex},t) \geq 0$. As a result, we defined the boundary $x_0$ between the cortex and the cytoplasm as the position for which the temporal mean of the intensity variation is zero : $\langle \Delta i(x_0,t) \rangle_t = 0$ (Fig S1f).

[4] This average corresponds to the average of the black curves in Fig S1d for all $t < t_{act}$

We assumed that the peak of fluorescence emanating from the cortex is approximately symmetrical. For simplicity, the outer edge of the cell $x_{\text{out}}$ is then defined using the relationship: $x_{\max} - x_0 = x_{\text{out}} - x_{\max}$. Therefore, the fluorescence intensity changes in the cytoplasm and cortex are expressed as : $\Delta i_{\text{cyto}}(t) = \langle \Delta i(x_{\text{cyto}}, t) \rangle_{x_{\text{cyto}}}$ where the average is taken over $x_{\text{cyto}} \in [5\text{pix}, x_0]$, and $\Delta i_{\text{cortex}}(t) = \langle \Delta i(x_{\text{cortex}}, t) \rangle_{x_{\text{cortex}}}$, where the average is taken over $x_{\text{cortex}} \in [x_0, x_{\text{out}}]$.

Fluorescence intensity in an image pixel represents a proxy for fluorophore concentration; a 3-dimensional concentration for the cytoplasm and a 2-dimensional concentration for the cortex. We introduce a variable $\Delta I^{2D}_{\text{cortex}}$ which is proportional to the two-dimensional concentration in the cortex, and a variable $\Delta I^{3D}_{\text{cyto}}$ proportional to the three-dimensional concentration in the cytoplasm:

$$\Delta I^{2D}_{\text{cortex}}(t) = \langle \Delta I_{cell} \rangle_{t=0^-} \cdot h_{\text{cortex}} \cdot \Delta i_{\text{cortex}}(t) \quad \text{and} \quad \Delta I^{3D}_{\text{cyto}}(t) = \langle \Delta I_{cell} \rangle_{t=0^-} \cdot \Delta i_{\text{cyto}}(t) \tag{3}$$

where $h_{\text{cortex}}$ is defined as $x_{\text{out}} - x_0$. Here, the multiplication by $\langle \Delta I_{cell} \rangle_{t=0^-}$, where $t = 0^-$ corresponds to a time before activation, allows to average across cells with different levels of CRY2 expression, and to compare the different optogenetic measurements performed on the same cell.

In **Fig 3 and Fig S1g**, we plot $\Delta I^{2D}_{\text{cortex}}(t)$, which we consider to be proportional to the **membranous** [**CRY2**$^*$**−CIBN**], the surface density of CRY2-CIBN complexes recruited to the cortex at time t.

Additionally, in **Figs 4-6, Fig S1h and Fig S2d-f**, we define the proportion of CRY2 at the membrane relative to total CRY2 in the cell as:

$$R^* = \frac{S \Delta I^{2D}_{\text{cortex}}}{V \langle \Delta I_{cell} \rangle_{t=0^-}} = \frac{S h_{\text{cortex}}}{V} \Delta i_{\text{cortex}}(t), \tag{4}$$

a dimensionless quantity, where $S$ is the cell surface and $V$ is the cell volume. Here, we assume that the value $V \langle \Delta I_{cell} \rangle (t)$ represents the total amount of CRY2 available for activation at time $t$, and $S h_{\text{cortex}}$ is the volume of the cortex.

Similarly, we quantify the recruitment of myosin MRLC-iRFP to the cortex with:

$$R^*_{\text{myosin}} = \frac{S \Delta I^{2D}_{\text{myosin,cortex}}}{V \langle \Delta I_{\text{myosin,cyto}} \rangle_{t=0^-}} \tag{5}$$

where various quantities are defined in the same way as for CRY2.

#### 1.3 Fluorescence intensity profile in local activation experiments

To analyse experiments involving optogenetic activation localised to a small region at the cell boundary, we first detected the cell contours in time from images of cortical myosin-iRFP using the Fiji plugin "JFilament" [13]. We then adapted our image analysis pipeline to measure fluorescence intensity as a function of the angular position $\theta$ relative to the activation site $\theta_{\text{act}}$. The angle $\theta$ and activation angle $\theta_{\text{act}}$ was computed as $atan2(y - y_{\text{center}}, x - x_{\text{center}})$, where ($x$, $y$) are the coordinates of a given point on the contour, and ($x_{\text{center}}$, $y_{\text{center}}$) corresponds to the centroid of the cell. The same segmentation and normalization steps were applied as described previously, but instead of averaging over the entire cortex, we extracted $I_{\text{cortex}}(\theta, t)$ along the contour, averaging over the cortical thickness $h_{\text{cortex}}$.

An arc length coordinate on the contour, $-L \le s \le L$, was also computed, using the matlab function arclength with the linear method [6]. Here, $L$ is half of the total contour length and we define the arc length such that $s(\theta_{act}) = 0$ where $\theta_{act}$ is the angle of the activation point relative to the center of the cell. To compare fluorescence profiles between different cells, we also defined a normalised arc length $\bar{s} = s/L$ with $-1 \le \bar{s} \le 1$.

An example of the change in normalised fluorescence as a function of both $\theta - \theta_{\text{act}}$ and $s$ is shown in Fig. 2d and Fig. 2e, respectively. The spatiotemporal recruitment dynamics of [CRY2$^*$ − CIBN] can be visualised in a 3D plot of the change in normalised CRY2 fluorescence as a function of $\theta - \theta_{\text{act}}$ and time; an example is shown in Fig. 2f.

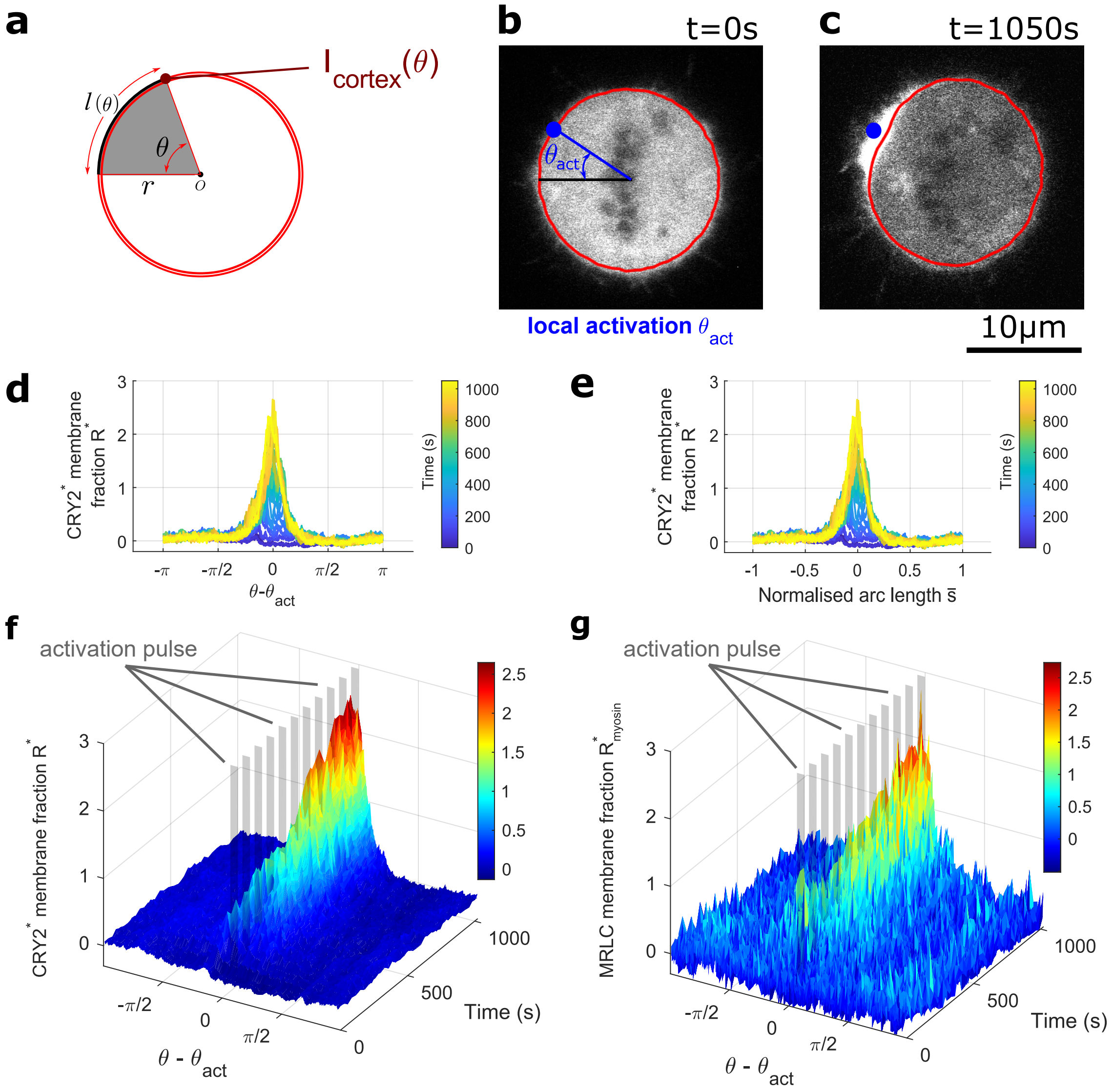


Supplementary Figure 2: **(a)** Diagram illustrating the geometry used to define the angular position $\theta$, the arc length $l$, and the cortical intensity $I_{\text{cortex}}$. The point $O$ corresponds to the cell centroid, with coordinates $(x_{\text{center}}, y_{\text{center}})$. **(b)** Image of CRY2-mCherry localisation in the cell at $t = 0$, before the first activation. The activation occurs in a small region (blue) located on the cell boundary at $\theta_{\text{act}}$. **(c)** Same as (b), but at $t = 1050$ s, after multiple exposures of the activation region (blue) to 488nm light. **(d)** Change in the CRY2* membrane fraction at different times after activation for different angular positions parameterised as $\theta - \theta_{\text{act}}$, with color representing time. **(e)** Same as (d), but plotted as a function of the normalised arc length $\bar{s}$. **(f)** 3D plot of the change in the CRY2* membrane fraction as a function of $\theta - \theta_{\text{act}}$ and time, showing the spatial and temporal evolution of recruitment. The grey bars represent spatio-temporal optogenetic pulses. **(g)** 3D plot of the change in the MRLC membrane fraction as a function of $\theta - \theta_{\text{act}}$ and time, showing the spatial and temporal evolution of recruitment.The grey bars represent spatio-temporal optogenetic pulses.

#### 2 Model of the optogenetic system

##### 2.1 Binding kinetics of CRY2 and CIBN following exposure to light

Here we describe a simple model for the binding kinetics of CRY and CIBN. Activated CRY2 is denoted CRY2*. We assume that only CRY2* can bind to CIBN, consistent with reports of low dark state binding between the two proteins [9, 15], with binding rate $k^+$. CRY2* spontaneously inactivates with rate $k$, with the same rate when the molecule is free or bound to CIBN. Furthermore, we assume that CRY2* detaches from CIBN only when it returns to its inactive state CRY2, and that the dissociation of the CRY2* − CIBN complex is immediate when CRY2 becomes inactive. This set of assumptions correspond to the following chemical reactions:

$$[\mathrm{CIBN}]^{2\mathrm{D}} + [\mathrm{CRY2}^*] \xrightarrow{k_+} [\mathrm{CRY2}^* - \mathrm{CIBN}]^{2\mathrm{D}} \xrightarrow{k} [\mathrm{CIBN}]^{2\mathrm{D}} + [\mathrm{CRY2}]\ , \tag{6}$$

$$[\mathrm{CRY2}^*] \xrightarrow{k} [\mathrm{CRY2}]\ , \tag{7}$$

where $[\mathrm{CRY2}^*]$, $[\mathrm{CRY2}]$ are the concentrations of active and inactive CRY2 in the cytoplasm, and $[\mathrm{CIBN}]^{2\mathrm{D}}$, $[\mathrm{CRY2}^* - \mathrm{CIBN}]^{2\mathrm{D}}$ are the two-dimensional concentrations of CIBN and CRY2*-CIBN at the cell surface.

The change of concentrations of CRY2* and CRY2*-CIBN associated to this set of chemical reactions can be described by the following differential equations:

$$\frac{d[\mathrm{CRY2}^*]}{dt} = -k[\mathrm{CRY2}^*] - k_+\frac{S}{V}[\mathrm{CRY2}^*][\mathrm{CIBN}]^{2\mathrm{D}}\ , \tag{8}$$

$$\frac{d[\mathrm{CRY2}^* - \mathrm{CIBN}]^{2\mathrm{D}}}{dt} = k_+[\mathrm{CRY2}^*][\mathrm{CIBN}]^{2\mathrm{D}} - k[\mathrm{CRY2}^* - \mathrm{CIBN}]^{2\mathrm{D}}\ , \tag{9}$$

where $V$ is the cell volume and $S$ the cell surface area.

Because of conservation of membrane-bound CIBN and of CRY2:

$$[\mathrm{CRY2}^* - \mathrm{CIBN}]^{2\mathrm{D}} + [\mathrm{CIBN}]^{2\mathrm{D}} = [\mathrm{CIBN}]^{2\mathrm{D}}_{\mathrm{tot}} \tag{10}$$

$$\frac{S}{V}[\mathrm{CRY2}^* - \mathrm{CIBN}]^{2\mathrm{D}} + [\mathrm{CRY2}^*] + [\mathrm{CRY2}] = [\mathrm{CRY2}]_{\mathrm{tot}}\ , \tag{11}$$

hence the concentration equations can be rewritten:

$$\frac{d[\mathrm{CRY2}^*]}{dt} = -k[\mathrm{CRY2}^*] - k_+\frac{S}{V}[\mathrm{CRY2}^*]([\mathrm{CIBN}]^{2\mathrm{D}}_{\mathrm{tot}} - [\mathrm{CRY2}^* - \mathrm{CIBN}]^{2\mathrm{D}})\ , \tag{12}$$

$$\frac{d[\mathrm{CRY2}^* - \mathrm{CIBN}]^{2\mathrm{D}}}{dt} = k_+[\mathrm{CRY2}^*][\mathrm{CIBN}]^{2\mathrm{D}}_{\mathrm{tot}} - (k_+[\mathrm{CRY2}^*] + k)[\mathrm{CRY2}^* - \mathrm{CIBN}]^{2\mathrm{D}}\ . \tag{13}$$

The last equation can also be rewritten in a convenient form:

$$\frac{d}{dt}\frac{[\mathrm{CRY2}^* - \mathrm{CIBN}]^{2\mathrm{D}}}{[\mathrm{CIBN}]^{2\mathrm{D}}_{\mathrm{tot}}} = k_+[\mathrm{CRY2}^*] - (k_+[\mathrm{CRY2}^*] + k)\frac{[\mathrm{CRY2}^* - \mathrm{CIBN}]^{2\mathrm{D}}}{[\mathrm{CIBN}]^{2\mathrm{D}}_{\mathrm{tot}}} \tag{14}$$

where $\frac{[\mathrm{CRY2}^* - \mathrm{CIBN}]^{2\mathrm{D}}}{[\mathrm{CIBN}]^{2\mathrm{D}}_{\mathrm{tot}}}$ is the fraction of bound CIBN.

Assuming $[\mathrm{CRY2} - \mathrm{CIBN}] \ll [\mathrm{CIBN}]_{\mathrm{tot}}$, one can simplify the two equations into two linear equations:

$$\frac{d[\mathrm{CRY2}^*]}{dt} = -k[\mathrm{CRY2}^*] - k_+[\mathrm{CIBN}]^{2\mathrm{D}}_{\mathrm{tot}}\frac{S}{V}[\mathrm{CRY2}^*] \tag{15}$$

$$\frac{d[\mathrm{CRY2}^* - \mathrm{CIBN}]^{2\mathrm{D}}}{dt} = k_+[\mathrm{CIBN}]^{2\mathrm{D}}_{\mathrm{tot}}[\mathrm{CRY2}^*] - k[\mathrm{CRY2}^* - \mathrm{CIBN}]^{2\mathrm{D}} \tag{16}$$

We experimentally verified that this condition is fulfilled by comparing the quantity of CIBN-EGFP-CAAX mRNA transcripts to CRY2-mCherry mRNA transcripts using qRT-PCR for all of the cell lines in this study (Fig S7a); assuming that for these proteins, the amount of protein is proportional to the amount of mRNA. These measurements indicate that the amount of mRNA transcript for CIBN-EGFP-CAAX is on average 1000-fold larger than CRY2-mCherry mRNA transcript within the cell population, although this varies from cell to cell.

These equations can be solved analytically with the initial conditions: $[\mathrm{CRY2}^*](0) = [\mathrm{CRY2}^*]_0$ and $[\mathrm{CRY2}-\mathrm{CIBN}](0) = 0$. Indeed, an initial pulse of light sets the initial concentration of activated optogenetic protein $[\mathrm{CRY2}^*]_0$. By setting $\bar{k} = k + \frac{S}{V} k_+ [\mathrm{CIBN}]_{\mathrm{tot}}$:

$$[\mathrm{CRY2}^*](t) = [\mathrm{CRY2}^*]_0 \exp(-\bar{k}t) \tag{17}$$

$$[\mathrm{CRY2}^* - \mathrm{CIBN}]^{2\mathrm{D}}(t) = [\mathrm{CRY2}^*]_0 \frac{V}{S} \left(\exp(-kt) - \exp(-\bar{k}t)\right) \tag{18}$$

This functional form is fitted to experimental measurement of the CRY2 membrane fraction $R^*$ (Figs. 4-6) as defined in section 1.2, assuming that $R^*$ and $\frac{S[\mathrm{CRY2}^*-\mathrm{CIBN}]^{2\mathrm{D}}}{V[\mathrm{CRY2}]_{\mathrm{tot}}}$ are proportional :

$$\mathrm{R}^*(t) = R_0^* \left(\exp(-kt) - \exp(-\bar{k}t)\right) \tag{19}$$

with $R_0^* = \frac{[\mathrm{CRY2}^*]_0}{[\mathrm{CRY2}]_{\mathrm{tot}}}$. The term $V[\mathrm{CRY2}]_{\mathrm{tot}}$ represents the total amount of CRY2 available for activation. Consequently, $R_0^*$ represents the inital fraction of CRY2 that is activated.

#### 2.2 Conversion of CRY2 into CRY2$^*$ as a function of photonic energy

Here we discuss a simple description of the conversion rate of CRY2 into CRY2$^*$, as a function of the total laser energy. We assume that a fraction $\alpha$ of the CRY2 in the cell can actually be activated. We also assume that between $t$ and $t + dt$, the probability of activating a CRY2 is proportional to the energy delivered during that time, $p = \kappa P_{\mathrm{act}} dt$, where $P_{\mathrm{act}}$ is the laser power, $P_{\mathrm{act}} dt$ the energy delivered, and $\kappa$ a proportionality coefficient. We also assume that during the activation time $\tau_{\mathrm{act}}$, activation is irreversible. Then the average fraction of activated CRY2 during a total activation time $\tau_{\mathrm{act}}$ is given by:

$$R_0^*(P_{\mathrm{act}}\tau_{\mathrm{act}}) = \frac{[\mathrm{CRY2}^*]_0(P_{\mathrm{act}}\tau_{\mathrm{act}})}{[\mathrm{CRY2}]_{\mathrm{tot}}} = \alpha(1 - \exp(-\kappa P_{\mathrm{act}}\tau_{\mathrm{act}})) \tag{20}$$

and is a simple exponentially relaxing function of the total energy $E_{\mathrm{act}} = P_{\mathrm{act}}\tau_{\mathrm{act}}$.

Experimentally, such a scaling is observed when we vary $E_{act}$ by varying either $\tau_{act}$ or $P_{act}$ (**Fig 4d-e, l**). In our experiments, we measured an average $\overline{\alpha} = 0.48 \pm 0.12$.

#### 2.3 Delay model for cortical myosin and cortical tension

We denote $\Delta T$ the change in cortical tension (Supplementary Fig S4c) and $R^*_{\mathrm{myosin}}$ the change in myosin membrane fraction following a change in the concentration of the CRY2-mCherry-RhoGEF at the cortex. To predict the temporal evolution of these variables, we propose that this response is proportional to the change in membrane fraction of CRY2-mCherry-RhoGEF $R^*(t)$ but occurs with a delay. Therefore, we introduce the response coefficients $\kappa_{\mathrm{myosin}}$ (expressed in $\mathrm{s}^{-1}$) and $\kappa_{\mathrm{tension}}$ (expressed in $\mathrm{N} \cdot \mathrm{m}^{-1} \cdot \mathrm{s}^{-1}$), as well as the delays $\tau_{\mathrm{myosin}}$ and $\tau_{\mathrm{tension}}$ (both expressed in $s$).

$$\frac{dR^*_{\mathrm{myosin}}}{dt} = \kappa_{\mathrm{myosin}} R^*(t) - \frac{1}{\tau_{\mathrm{myosin}}} R^*_{\mathrm{myosin}}(t) \tag{21}$$

$$\frac{d\Delta T}{dt} = \kappa_{\mathrm{tension}} R^*(t) - \frac{1}{\tau_{\mathrm{tension}}} \Delta T(t) \tag{22}$$

Starting from the double exponential expression of $R^*(t)$ (Eq. 19), we obtain analytical expressions for $R^*_{\mathrm{myosin}}(t)$ and $\Delta T(t)$. For both quantities, the differential equation takes the following form:

$$\frac{dy}{dt} = \kappa R_0^* \left(e^{-kt} - e^{-\bar{k}t}\right) - \frac{1}{\tau} y \tag{23}$$

where $y(t)$ and $\kappa$ denote either $R_{\text{myosin}}(t)$ and $\kappa_{\text{myosin}}$, or $\Delta T(t)$ and $\kappa_{\text{tension}}$. The solution for $y(t)$ with the initial condition $y(0) = 0$ is given by:

$$y(t) = R_0^* \frac{\kappa\tau e^{-\frac{t}{\tau}}}{(1-k\tau)(1-\bar{k}\tau)} \left[ k\tau \left( e^{(1-\bar{k}\tau)\frac{t}{\tau}} - 1 \right) - \bar{k}\tau e^{(1-k\tau)\frac{t}{\tau}} + e^{(1-k\tau)\frac{t}{\tau}} - e^{(1-\bar{k}\tau)\frac{t}{\tau}} + \bar{k}\tau \right] \quad (24)$$

#### 2.4 Fitting Procedures

All fitting was performed using the MATLAB function `lsqcurvefit`. For fits on averaged data (Fig. 2a-c; Fig. 3d-e; Fig. 5d-f), the fitting was first conducted on the CRY2 fluorescence signal used in each figure (Fig. 2a-c : $I_{cortex}/I_{cyto}$; Fig. 3d-e : $\Delta I^{2D}_{cortex}$; Fig. 5d-f :$R^*$) to extract the parameters $k$, $\bar{k}$, and $[\text{CRY2}^*]_0$. These values were then fixed for the subsequent fits of the temporal evolution of $\Delta T$ (tension variation) and myosin recruitment signal to determine the delay times $\tau$ and response coefficients $\kappa$ (i.e. fits for Fig. 3d-e, and Fig. 5d-f).

For fits on data from individual cells (i.e. box plots in Fig. 2a-c insets; Fig. 3f-h; and Fig. S6i-n), the fitting was performed similarly, starting with the CRY2 fluorescence signal for each figure (Fig. 2a-c insets: $I_{cortex}/I_{cyto}$; Fig. 3f-h and Fig. S6i-n: $\Delta I^{2D}_{cortex}$) was first fitted to obtain cell-specific values of $k$, $\bar{k}$, and $[\text{CRY2}^*]_0$. The mean values of $\langle k \rangle$ and $\langle \bar{k} \rangle$, together with their standard deviation (SD), were then computed across cells. Fits of $\Delta T$ were performed using these averaged parameters constrained within ranges of $\langle k \rangle \pm \text{SD}$ and $\langle \bar{k} \rangle \pm \text{SD}$.

In cases where data sets were acquired from the same cell with different activation times (Figs 4 and 5), we assumed that the parameters $k$ and $\bar{k}$ do not change with activation time and are therefore shared across the dataset. Consequently, all curves acquired on the same cell (Fig. 4d-l) or all averaged curves (Fig. 5d-f) were fitted simultaneously using a common set of parameters $k$ and $\bar{k}$.

### 3 Active surface model for the cell cortex

To predict the deformation of a cell upon localised optogenetic actuation, we first formulate a mechanical model of the cellular actomyosin cortex as an active viscous surface [11, 12, 2]. This 2-dimensional representation is justified because the cortical thickness ($h \approx 200nm$, [4]) is much smaller than the cell radii ($R \approx 10\mu m$, see Table 2). We further assume that bending rigidity is negligible. The cortical thickness is taken to be constant throughout, due to the rapid turnover of cortical components (time scale of $\sim 1 - 100s$ [8]) relative to the deformation time scale of $\sim 10$ minutes (see main text Fig. 6 b).

We denote $g^{ij}$ the metric tensor and $C^{ij}$ the curvature tensor of a curved surface representing the actomyosin cortex and the cell membrane. We refer to Refs [12, 10] for further notations of differential geometry. The constitutive equation for the tension tensor of the active viscous cell cortex is taken as

$$t^{ij} = (\gamma + (\eta_b - \eta)v^k_k)g^{ij} + 2\eta v^{ij}, \quad (25)$$

where $\eta$ and $\eta_b$ are the 2-dimensional shear and bulk viscosities (we set $\eta = \eta_b$ for simplicity), and $\gamma$ is an isotropic active tension. Spatial gradients in $\gamma$ are then driving flows and deformations of this surface. The strain rate tensor $v^{ij}$ is given by [12]

$$v^{ij} = \frac{1}{2}(\nabla^i v^j + \nabla^j v^i) + C^{ij} v_n, \quad (26)$$

with $\mathbf{v}$ the surface velocity, which has components tangential $(v_i, v_j)$ and normal $(v_n)$ to the surface.

The mechanical equilibrium of a fluid surface with material properties defined by equation (25) is given by the local force balance equations. Projected on the tangential and normal directions, those read

$$\nabla_i t^{ij} = 0 \quad (27)$$
$$C_{ij} t^{ij} = P. \quad (28)$$

Here, we have introduced a hydrostatic uniform pressure $P$ acting on the cortical surface from the inside, resulting from the cytoplasmic fluid enclosed by the cell membrane.

An optogenetic actuation localised to a small region of the cortex defines an axis of symmetry for the cellular shape (see main text Fig. 6 c). To capture the resulting axisymmetric deformation, the cell shape $\mathbf{X} \in \mathbb{R}^3$ is parametrised as

$$\mathbf{X}(\phi, s) = (x(s)\cos\phi, x(s)\sin\phi, z(s)), \tag{29}$$

with the arc length coordinate $s \in [0, L]$ defined along the cell outline from the actuation point (south pole) to the opposite cellular pole (north pole), i.e. $L$ is half of the cell contour length, and the angle of rotation $\phi \in [0, 2\pi]$ around the $z$-axis (see main text Fig. 6 c). We work in the Lagrangian framework to evolve the cell shape in time, in which material points move according to the full velocity vector $\mathbf{v}$,

$$\partial_t \mathbf{X} = \mathbf{v}. \tag{30}$$

In the axisymmetric parametrisation, the velocity is $\mathbf{v} = v^s \mathbf{e}_s + v_n \mathbf{n}$ and the two components are determined as instantaneous solutions of the force balance equations (27) and (28) on a given shape $\mathbf{X}(t)$. For details on the surface parametrisation, the parametrised form of equations (25)-(28) and the numerical solution of the resulting system of diffential equations see Khoromskaia and Salbreux [10].

#### 3.1 Dynamics of the tension profile

When optogenetic actuation happens in the centre of a cell, we observe an increase of the overall cortical tension (see main text Fig. 5). Therefore, if actuation happens locally at the cortex, we expect a local increase of tension and a spatial pattern $\gamma(s, t)$ will develop over time, which will determine the cellular shape change.

We thus take the tension profile to consist of a constant tension $\gamma_0$ prior to optogenetic activation at time $t = t_{act}$ and a spatial profile $\gamma_{act}(s, t)$ that builds up after activation,

$$\gamma(s, t) = \gamma_0 + \gamma_{act}(s, t). \tag{31}$$

Following the dynamical response of tension derived for fully activated cells (Eq. 1 in the main text and 22 of this document), the time evolution of the induced tension can be written as

$$\partial_t \gamma_{act}(s, t) = \kappa_{\text{tension}} \bar{R}^*(s, t) - \frac{1}{\tau_{\text{tension}}} \gamma_{act}(s, t). \tag{32}$$

Here, $R^*(s, t)$ is the spatial profile of the proportion of cortex-bound activated CRY2*, as defined in Eq. 4, and $\bar{R}^*(s, t)$ is the von Mises fit of this profile as described in Sec. 3.2.

Numerically, the Lagrangian time update of the full tension profile is performed by

$$\gamma(s_{new}, t + dt) = \gamma(s, t) + dt \left( \kappa_{\text{tension}} \bar{R}^*(s, t) - \frac{1}{\tau_{\text{tension}}} (\gamma(s, t) - \gamma_0) \right) \tag{33}$$

where $s$ and $s_{new}$ are the arc length grids at time $t$ and $t + dt$, respectively (see [10] for details). In this way, we take into account that the dynamical response is associated to material points (i.e. parts of the cortex) that are moving with the flow. Here we have not included a possible effect of dilution or enrichment associated to non-zero flow divergence (i.e. a term $-\gamma v_k^k$ on the right hand side of the dynamical equation).

Additionally, we calculate the optogenetically induced myosin profile at the cortex as

$$\partial_t c_{\text{myo}}(s, t) = \kappa_{\text{myosin}} \bar{R}^*(s, t) - \frac{1}{\tau_{\text{myosin}}} c_{\text{myo}}(s, t), \tag{34}$$

which moves with the material points analogously to Eq.(33).

#### 3.2 Input from experimental profiles

Normalised profiles of CRY2-mCherry increase are extracted from the fluorescence signal along the cell contour as described in Section 1.3. In order to produce a model input, these profiles are fitted with a smooth function having the required boundary conditions. At each time of experimental measurement $t_i \in \{0s, 30s, 90s, ..., t_{max}\}$, the profile $R^*(\bar{s}, t_i)$ on the normalised arc length $\bar{s} = s/L$ (see main text Fig. 6e) is fitted with a von Mises function in the form

$$f(\bar{s}|a, b, k) = a \exp(k \cos(\pi(\bar{s} - b))) \tag{35}$$

with fitting constants $a, b, k \in \mathbb{R}$, $a, k > 0$. We choose this functional representation of the data, as the shape is similar to a Gaussian, but fulfills the boundary conditions $\partial_{\bar{s}} f = 0$ at the poles of the cell for $b = 0$, which are required by axisymmetry. Due to small deviation of experimentally observed cell shapes from ideal axisymmetry, the fitted peak may not be exactly centred at $\bar{s} = 0$. Therefore, b is introduced as a fitting parameter. For the function (35) the height of the peak is given by $a \exp(k)$ and the full width at half maximum (FWHM) is given by

$$\text{FWHM} = \frac{2}{\pi} \arccos\left(1 - \frac{1}{k} \ln 2\right). \tag{36}$$

These quantities are shown in main text (Fig. 6e) for one example cell (Cell 1). To obtain values of the parameters $a(t)$ and $k(t)$ for all $t \in [0, t_{max}]$, the values at $t_i$ are interpolated piece-wise linearly. In simulations we set $b = 0$ throughout, i.e. the peak of CRY2-mCherry is localised directly at the south pole of the cell.

We reconstruct the cell volume as an integral of the cell outline assuming axisymmetry, $V_{cell} = \pi \int_0^{z_{max}} x^2 dz$. This is done separately for each half of the cell outline and then averaged, as the cell shape is not perfectly axisymmetric. The cell volume changes non-monotonically by up to 10% as the cells are flattening at one pole (see main Fig. 6 b), we therefore decided to enforce the observed volume change in the simulations. We obtain $V_{cell}(t)$ as a piece-wise linear interpolant of the measured $V_{cell}(t_i)$. The pressure $P(t)$ then acts as a Lagrange multiplier to satisfy the integral constraint $\partial_t V = 2\pi \int ds\, x v_n = (V_{cell}(t + dt) - V_{cell}(t))/dt$, when solving for the velocity field at time $t$ (see also [10]).

#### 3.3 Choice of parameters and results of numerical simulation

Simulations are initialised at time $t = 0$ with spherical cells of radius $R_c$, with physical parameters as given in Tables 1 and 2.

We base our reference viscosity $\eta_0$ on viscosity values reported previously for mitotic cells ($3.25\times10^3\text{Pa}\cdot\mu\text{m}\cdot\text{s}$ [7]) and migrating cells ($2.7\times10^3\text{Pa}\cdot\mu\text{m}\cdot\text{s}$ [1]) and chose $\eta_0 = 3\times10^3\text{Pa}\cdot\mu\text{m}\cdot\text{s}$ as a reference value. The viscosity is then gradually increased to capture the time scale of deformation for Cell 1 (see Suppl. Fig. 3 a). Simulation with $\eta = \eta_0$ results in profiles of tension $\gamma(s, t)$ and of myosin $c_{\text{myo}}(s, t)$ which develop a cusp at the south pole ($c_{\text{myo}}$ profile at the corresponding time point is shown in Suppl. Fig. 3 b), possibly due to large tangential cortical flows. Viscosities of $\eta = 10\eta_0, 100\eta_0$ show realistic tension and myosin profiles over the whole time range. Specifically, $\eta = 10\eta_0$ also reproduces the experimental speed of deformation, therefore this value is used for all remaining simulations, including those for Cell 2 and Cell 3. From the time scale of deformation of $\tau \approx 200s$, which is similar across all three cells (see main text Fig. 6 b), and the reference tension $\gamma_0 = 0.5\text{mN}\,\text{m}^{-1}$, we can roughly estimate the viscosity as $\eta \sim \gamma_0\tau \approx 10^5\text{Pa}\cdot\mu\text{m}\cdot\text{s}$.

The parameters $\tau_{\text{tension}}$ and $\kappa_{\text{tension}}$ are first measured for fully activated cells (see main text Fig. 5) and given in Table 1. However, to obtain in simulations the cellular deformation as observed experimentally, values of $\kappa_{\text{tension}}$ much smaller than $0.12\,\text{mN}\,\text{m}^{-1}\,\text{s}^{-1}$ were required for all three cells (see Suppl. Fig. 3 c,d,e). This led us to hypothesize that there could be a saturation effect at high values of membraneous fraction of CRY2, such that the linear relationship between tension and amount of CRY2 activation obtained from full cell activation experiments (Fig. 5i) no longer holds. For each cell, we systematically increased the proportionality factor, starting from

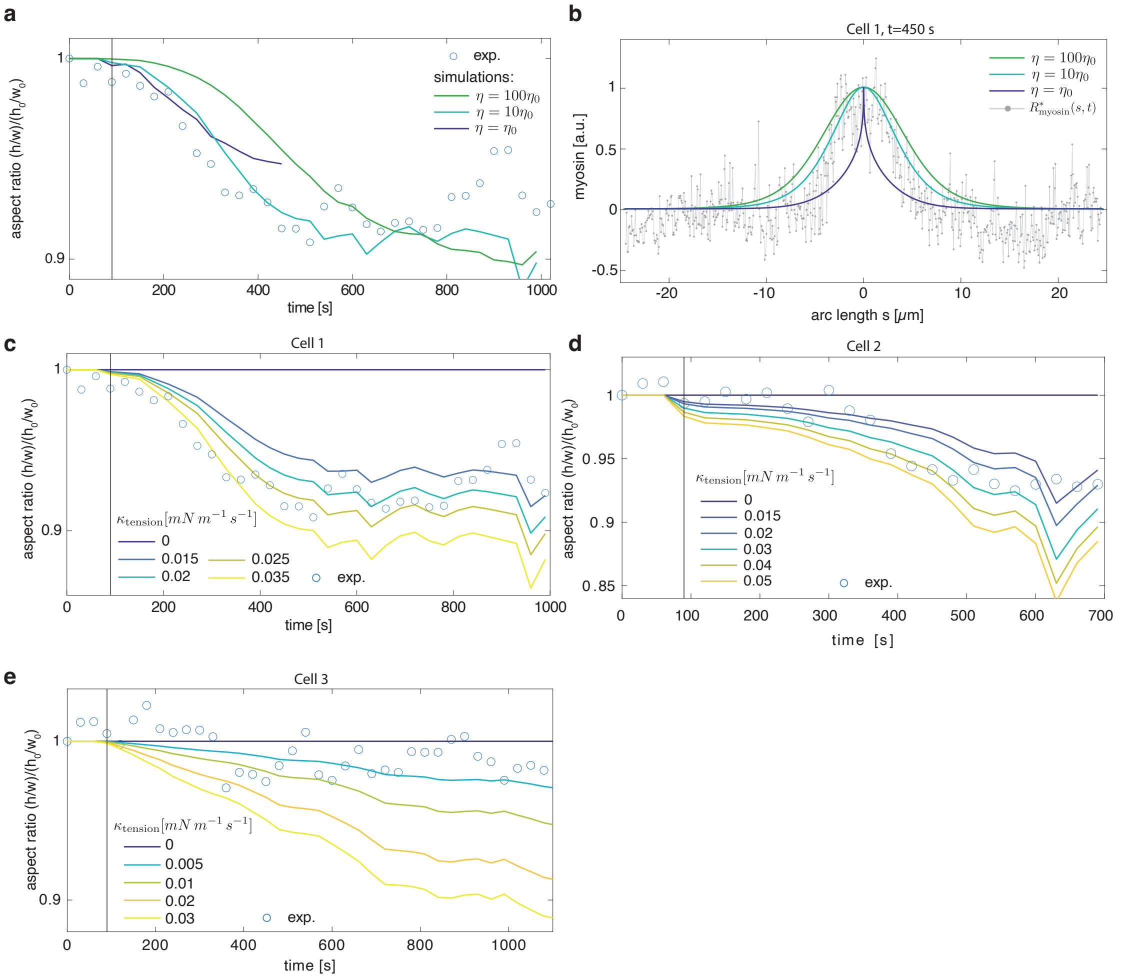


Supplementary Figure 3: **(a)** Time evolution of cell aspect ratio for Cell 1 from experiment and from simulations with $\kappa_{\mathrm{tension}} = 0.025\,\mathrm{mN\,m^{-1}\,s^{-1}}$ and different values of viscosity $\eta$. **(b)** Experimentally observed profile $R^*_{\mathrm{myosin}}$ at $t = 450$s (same data as Suppl. Fig. 2 e) and simulated induced myosin $c_{\mathrm{myo}}(s, t = 450s)$, from Eq.(34) with $\tau_{\mathrm{myosin}} = 120s$ (see main text Fig. 5 e), $\kappa_{\mathrm{myosin}} = 0.014$ (chosen to match the peak height of $R^*_{\mathrm{myosin}}$) and different values of viscosity $\eta$. **(c,d,e)** Cell aspect ratio from experiment and from simulations for different values of $\kappa_{\mathrm{tension}}$ for Cell 1, Cell 2, and Cell 3.

$\kappa_{\text{tension}} = 0$ (no deformation), to identify the value that minimises the error between the simulated and the experimental shape dynamics. The optimal values $\kappa_{\text{tension}} = \bar{\kappa}_{\text{tension}}$ are given in Table 3, together with the maximum tension increase at the location of activation in those simulations.

Other numerical parameters are chosen as follows: a fixed time step $dt = 10^{-3}$s, bending viscosity parameter $\eta_{cb} = 0.16\eta$, relative error $\varepsilon_{rel} = 10^{-3}$, absolute error $\varepsilon_{abs} = 10^{-6}$ (see [10] for details). Here, we do not make equations dimensionless for simulations, so time is given in seconds, lengths in $\mu$m and tension in mN $(\mu\text{m})^{-1}$.

| **parameter** | **value** | **description** |
|---|---|---|
| $\gamma_0$ | 0.5 $\text{mN}\,\text{m}^{-1}$ | reference tension (main text Fig. 5b) |
| $\eta$ | $10\eta_0 = 3 \times 10^4 \text{Pa} \cdot \mu\text{m} \cdot \text{s}$ | viscosity |
| $\tau_{\text{tension}}$ | 30 s | see main text (Fig. 5) |
| $\kappa_{\text{tension}}$ | $0.12\,\text{mN}\,\text{m}^{-1}\,\text{s}^{-1}$ | see main text (Fig. 5) |

Table 1: Default parameter values for active viscous surface model.

| **Cell** | **cell radius** $R_c$ | $\bar{\kappa}_{tension}$ $[\text{mN}\,\text{m}^{-1}\text{s}^{-1}]$ |
|---|---|---|
| Cell 1 | 7.94 µm | 0.025 |
| Cell 2 | 8.6 µm | 0.02 |
| Cell 3 | 9.67 µm | 0.005 |

Table 2: Overview of cell specific numerical parameters.

| **Cell** | $\max R^*$ | $\max \Delta\gamma$ $[\text{mN}\,\text{m}^{-1}]$ |
|---|---|---|
| Cell 1 | 2.13 | 2.48 |
| Cell 2 | 0.85 | 1.26 |
| Cell 3 | 1.4 | 0.21 |

Table 3: Overview of maximal membraneous CRY2 fraction ($\max R^*$) and maximal tension increase ($\max \Delta\gamma$) in the simulation for the different cells.

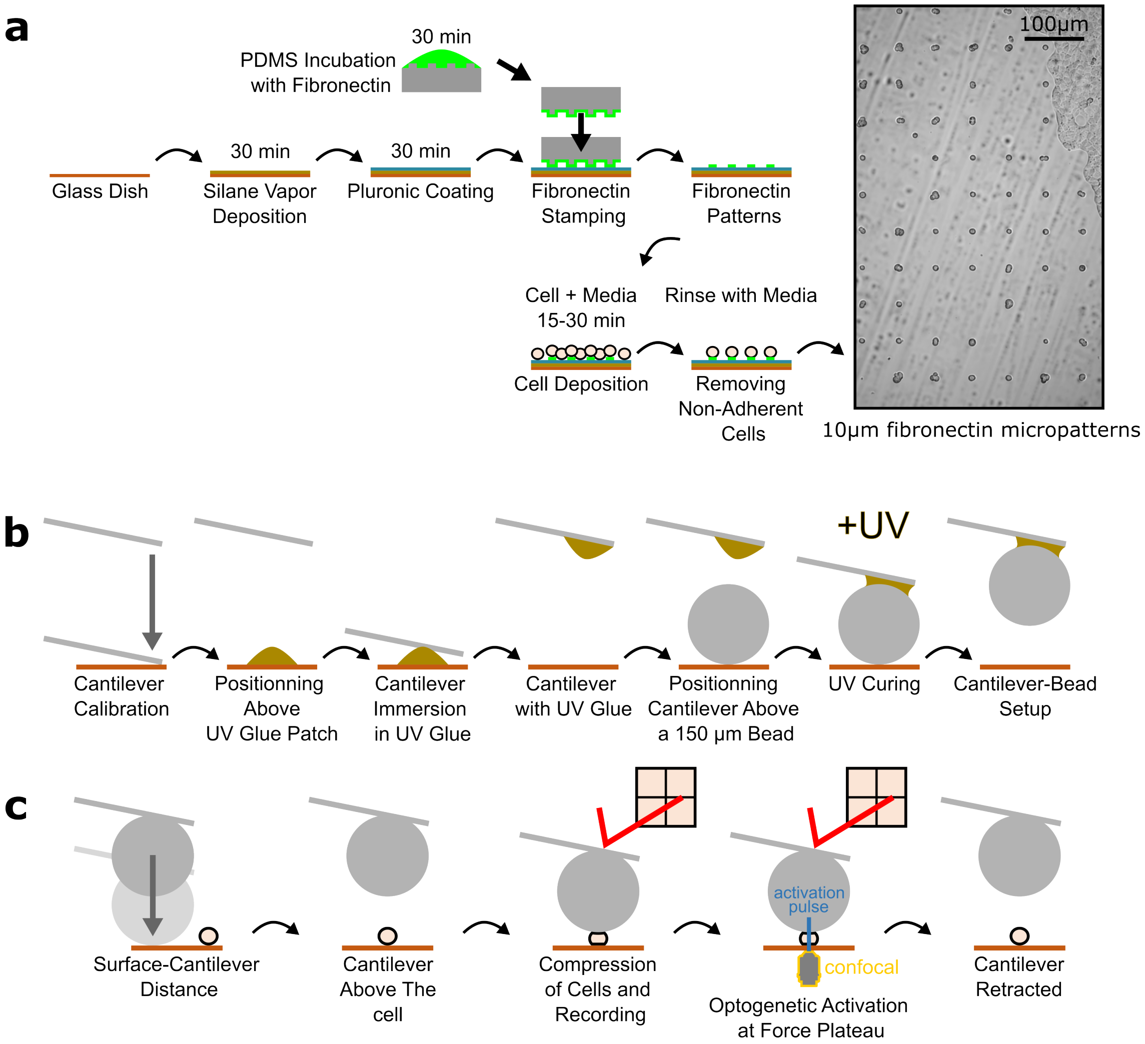


Supplementary Figure 4: **(a)** Diagrams illustrating the steps involved in creating fibronectin micropatterns using the PDMS stamping protocol. **(b)** Diagrams showing the process of attaching a 150 $\mu$m bead to a tipless cantilever. **(c)** Diagrams outlining the steps allowing measurement of cortical tension during the optogenetic experiment.

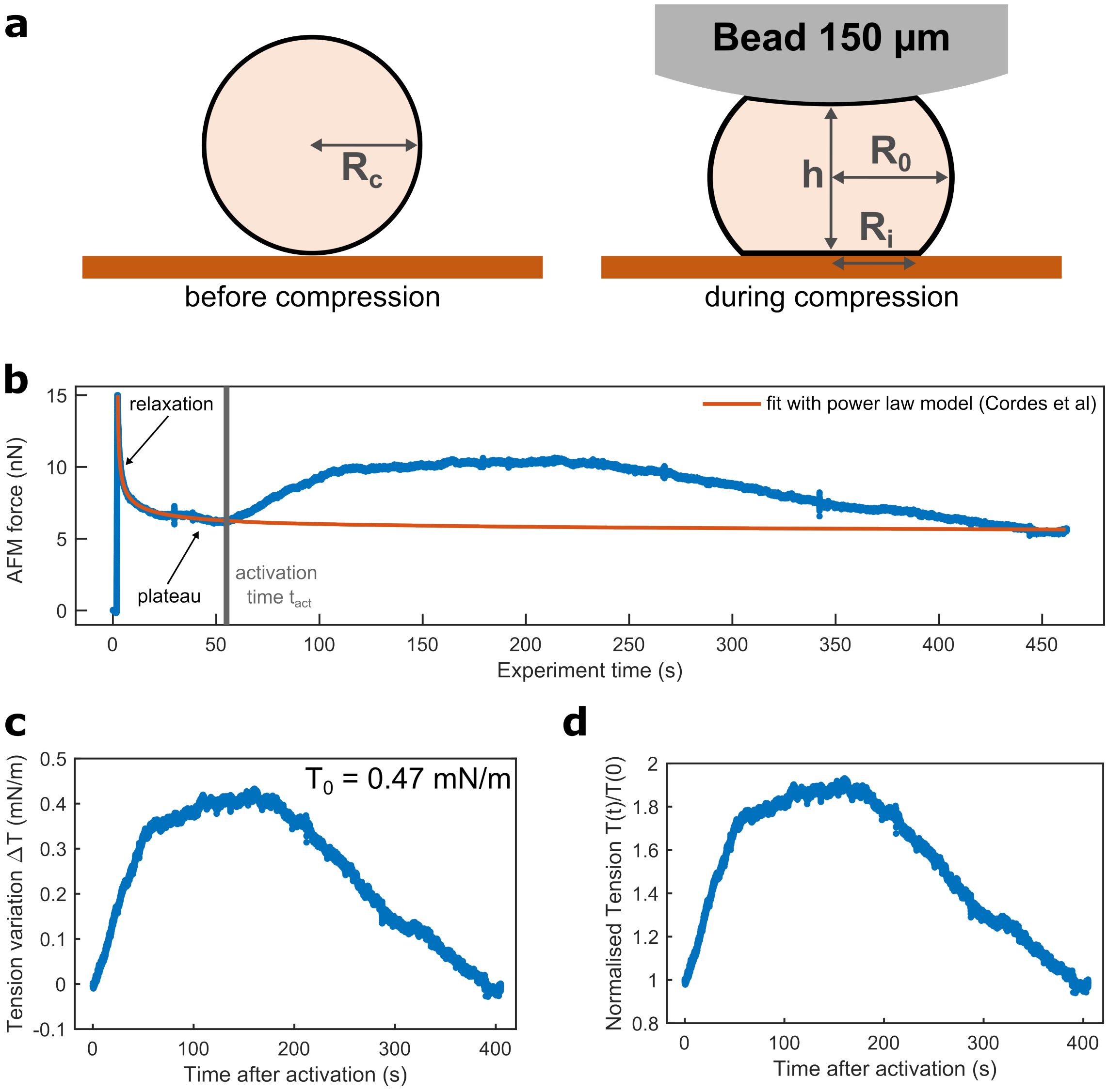


Supplementary Figure 5: **(a)** Diagram of a cell before (left) and during compression (right), showing the parameters: $R_c$, the radius of the cell before compression; $h$, the compression height, $R_0$, the radius at the equatorial plane during compression; and $R_i$, the radius of the contact surface. **(b)** Example of a force versus time measurement (blue curve) during cell compression with AFM, showing force in nN. The characteristic relaxation phase is followed by a plateau that allows determination of the basal tension $T_0 = 0.47$ mN/m. At $t_{\text{act}}$, the optogenetic system is activated, inducing an increase in the measured force. The orange curve represents a fit to the initial relaxation phase with a power-law model for the area compressibility modulus $K_A$ ([5]). **(c)** The tension variation $\Delta T(t)$ after optogenetic activation obtained from subtracting the blue curve in b from the orange curve. **(d)** The normalized tension after optogenetic activation, $T(t)/T_0 = (\Delta T + T_0)/T_0$.

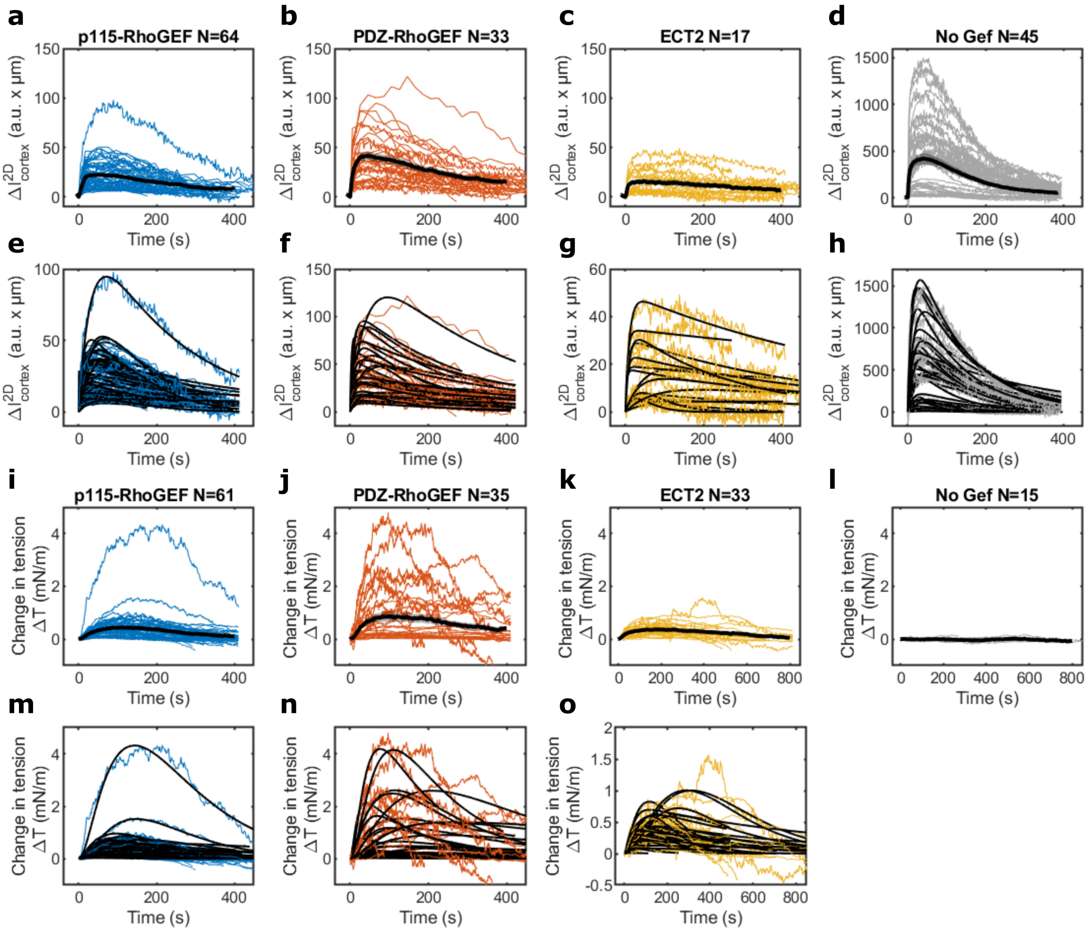


Supplementary Figure 6: Plots showing the temporal evolution of $\Delta I^{2D}_{cortex}$ considered to be the membranous [CRY2*-CIBN] (CRY2-mCherry-RhoGEF at the membrane) and the tension variation $\Delta T$ after optogenetic activation at $t = 0$ for different GEFs (p115-RhoGEF in blue, PDZ-RhoGEF in orange, ECT2 in yellow, and CRY2-mCherry without a RhoGEF domain in grey). The black curves in **(a-d)** and **(i-l)** represent the average values, with the grey regions indicating the standard error. The black curves in **(e-h)** and **(m-o)** represent fits to each experiment. For [CRY2*-CIBN] in (e-h), we fitted a double exponential model using Eq. 18. For the tension variation $\Delta T$ in **(m-o)**, we used the dealy model Eq. 24.

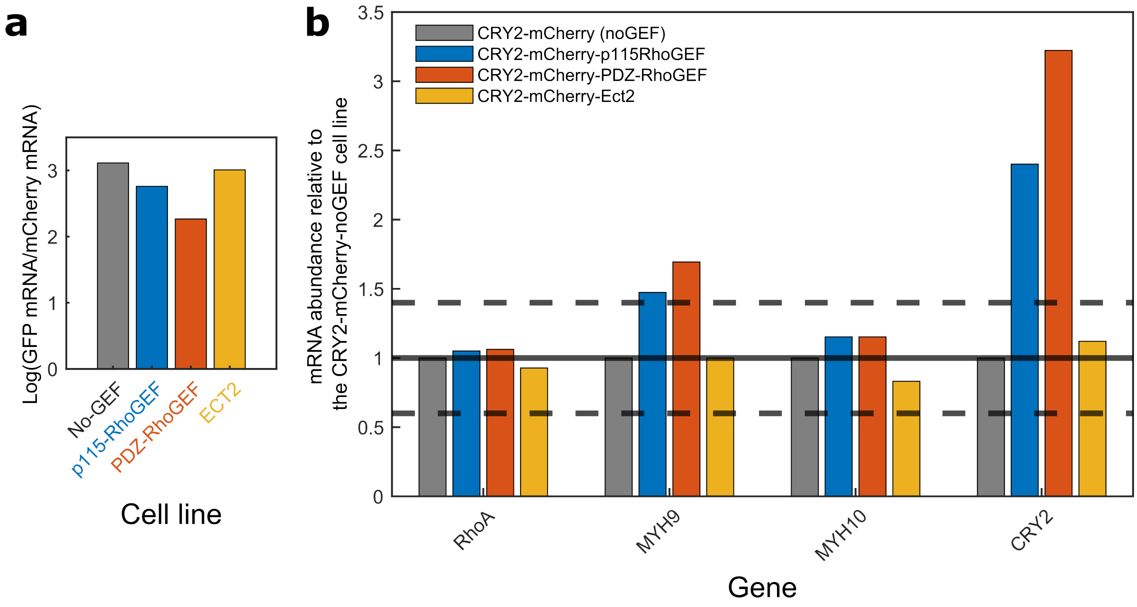


Supplementary Figure 7: **(a)** Bar chart showing the log ratio of GFP mRNA transcripts to mCherry mRNA transcripts for different cell lines: No GEF (grey), p115-RhoGEF (blue), PDZ-RhoGEF (orange), and Ect2 (yellow). **(b)** Bar chart showing the relative mRNA abundance normalized to the CRY2-mCherry-NoGEF cell line for the genes RhoA, MYH9, MYH10, and CRY2. Each gene is represented by four bars corresponding to the expression levels in No GEF (grey, control), p115-RhoGEF (blue), PDZ-RhoGEF (orange), and Ect2 (yellow) cell lines.

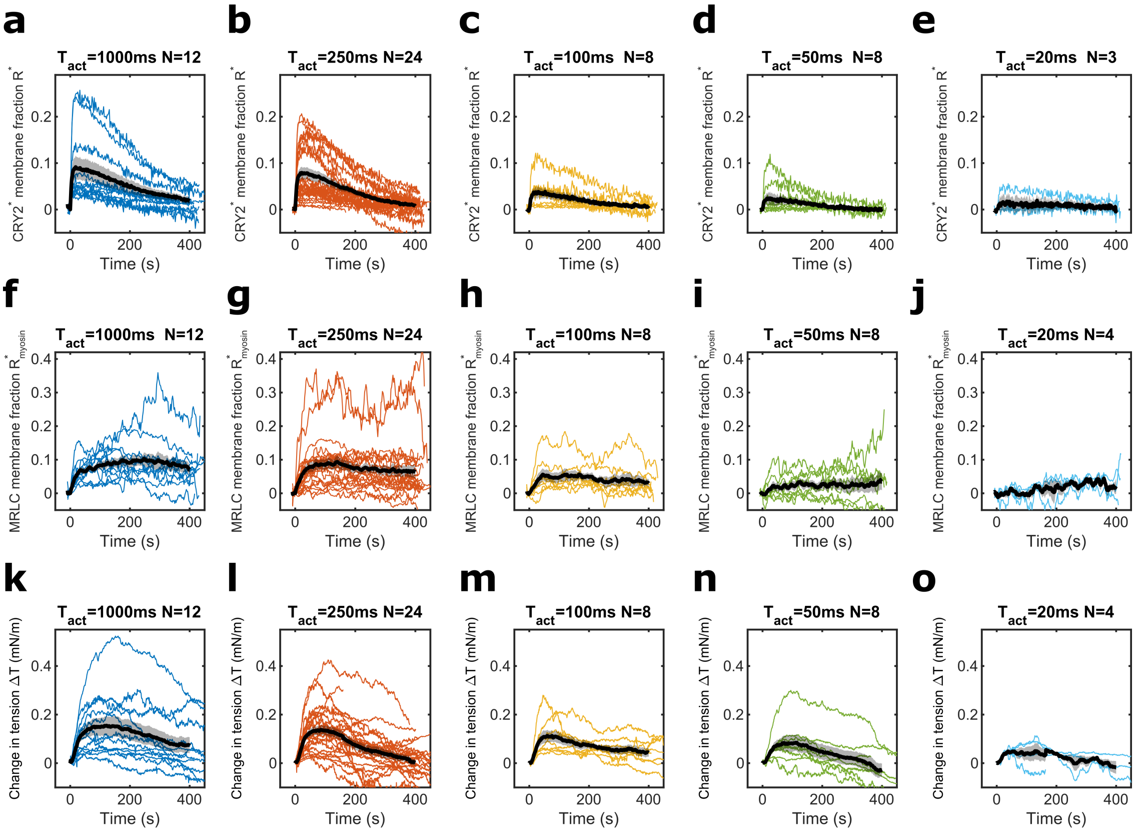


Supplementary Figure 8: **(a-e)** Plots showing the evolution of the CRY2* membrane fraction $R^*$ after optogenetic activation at $t = 0$ for the p115-RhoGEF cell line. **(f-j)** Plots showing the evolution of the MRLC membrane fraction $R^*_{myosin}$ after optogenetic activation at $t = 0$ for the p115-RhoGEF cell line. **(k-o)** Plots showing the evolution of the tension variation $\Delta T(t)$ after optogenetic activation at $t = 0$ for the p115-RhoGEF cell line. All plots show individual cells (colored curves) and the corresponding average (black curve) with the standard error shaded in grey. $N$ is the number of curves used for averaging. Data are presented for different activation times: **(a,f,k)** $t_{act} = 1000$ ms in blue, **(b,g,l)** $t_{act} = 250$ ms in orange, **(c,h,m)** $t_{act} = 100$ ms in yellow, **(d,i,n)** $t_{act} = 50$ ms in green, and **(e,j,o)** $t_{act} = 20$ ms in light blue.
